# Geo-climatic gradient shapes functional trait variations in *Salix eriocephala* Michx

**DOI:** 10.1101/057745

**Authors:** Arun S.K. Shunmugam, Raju Y. Soolanayakanahally, Robert D. Guy

**Affiliations:** Saskatoon Research and Development Centre, Agriculture and Agri-Food Canada, Saskatoon, Saskatchewan, S7N 0X2, Canada; Indian Head Research Farm, Agriculture and Agri-Food Canada, Indian Head, Saskatchewan,S0G 2K0, Canada; Department of Forest and Conservation Sciences, Faculty of Forestry, University of British,Columbia, Forest Sciences Centre, 2424 Main Mall, Vancouver, BC V6T 1Z4, Canada

**Keywords:** *Salix*, photosynthesis, stomatal density, phenology, mesophyll conductance

## Abstract

Intraspecific variations in seasonal phenology and growth physiology reflect adaptation to local climate. To explore the patterns of local adaptation along latitudinal and longitudinal clines, we used thirty-four populations of *Salix eriocephala* sourced from its natural ranges across Canada. The genotypes were examined for 6 phenology and 19 ecophysiology traits over two growing seasons under common garden condition. Photosynthetic assimilation rate (*A*) increased with increase in latitude when measured during free growth. In spite, the negative correlation between stomatal density and stomatal conductance (*g*_s_), higher *A* is facilitated *via* larger pore length among genotypes from short growing seasons. In addition, higher *A*, was positively associated with total leaf nitrogen and leaf mass per unit area. No population level differences were observed for water use-efficiency (Δ^13^C), however nitrogen isotope discrimination (δ^15^N) displayed latitudinal clines. Growing season phenological traits considered in this study accounted high heritability (*H*^2^=0.65-0.94). *Melampsora* rust infestation also displayed a strong latitudinal cline with high-latitude genotypes being more susceptible. Overall, the results support the hypothesis that functional trait variations are largely explained by climate of origin and facilitate selection of parents with superior adaptive traits in the Canadian willow improvement program for bioenergy and environmental applications.

## INTRODUCTION

The issue of local adaptation is a crucial problem in the view of climate change. Functional trait variation can provide insights into the physiological processes associated with a specie’s persistence across a range of environmental conditions resulting from local adaptation (Aitken and Whitlock 2013). Largely, local adaptation plays a significant role in maintaining genetic variation among plant populations (Hodkinson 1999), and to better understand the evolutionary mechanisms of local adaptation one needs to take into account the environmental factors that contribute to phenotypic variation in nature (Manzano-Piedras et al. 2014). Such divergent selection is tested using provenance trail or common garden approaches to better understand trait trade-off relationships (Stearns 1989). Hence, between-provenance variation probably represents the most powerful tool for testing hypotheses of climatic adaptation among perennial trees (Mátyás 1996), whereby environmental gradients have produced genetically based clinal patterns in phenotype through adaptive evolution.

Correlations between trait variation and geo-climatic factors may suggest the adaptive selection pressure exerted on a trait, thus demonstrating its adaptive significance. A negative latitudinal cline in tree height growth has been reported for deciduous (McKown et al. 2014a) and conifer seeds sourced from different provenances (Benomar et al. 2016). Although the height growth of trees is limited by growing conditions at high latitudes, it is not the case at low latitudes. Hence, the date of growth cessation is more important in differentiating among provenances height growth variations (Bridgewater 1990). Such trade-offs results from physiological and/or genetic links between traits and limit the possibility of evolution of beneficial traits (Weih 2003).

Starting with the classical work by Mooney and Billings (1961) on *Oxyra digyna*, studies have revealed population genetic differentiation in photosynthesis which follows latitudinal and/or elevational clines (Chapin and Oechel 1983, Robakowski et al. 2012). At the same time, Flood et al (2011) linked leaf morphological attributes (leaf thickness, stomatal densities, leaf nitrogen) influencing photosynthetic rates among ecotypes from diverse temperature and moisture regimes. Stomatal density and pore length determine maximum conductance of CO_2_ to thesite ofassimilation and also control transpirational water loss from leaves. Over the last 400-million years the stomatal design features have remained unchanged-i.e., to improve photosynthetic rates and enabling land plants to occupy vast geographic ranges with varying environments thus contributing to increased genetic diversity (Franks and Beerling 2009). A strong association among traits suggests that optimization of photosynthesis to local environment along a latitudinal gradient is one mechanism by which plant fitness is enhanced (Gornall and Guy 2007).

*Populus* species sampled over vast geographic ranges are well studied for their intraspecific variation in seasonal phenology and ecophysiology related traits (e.g., Soolanayakanahally et al. 2009, McKown et al. 2014a). However, this remains less investigated in native populations of *Salix* (willow) species adapted to varied growth habitats. For example, willows from varied habitats provided evidence for their differential expression of water usage strategies when subjected to drought stress under greenhouse conditions (Savage and Cavender-Bares 2011).The physiological, anatomical and biochemical processes driving genotypic variation in resource acquisition and use efficiencies are still elusive in willows. Δ^13^C is a time integrated proxy measure of intrinsic water-use efficiency (*WUE*_i_) and has been correlated to growth and biomass accumulation (Farquhar et al. 1989). Considerable genotypic variation in Δ^13^C has been reportedwithinthe genus *Populus* (Soolanayakanahally et al. 2009, Broeckx et al. 2014, McKown et al. 2014a) and in other tree species (Guy and Holowachuk 2001). These studies suggest that genetic variation in Δ^13^C is useful as a selection criterion for improved water-use efficiency. Observed genotypic differences in stomatal and mesophyll conductance to CO_2_ (*g_s_* and *g_m_*, respectively) were reported to affect Δ^13^C (Gresset et al. 2014, Barbour et al. 2015). In addition, leaf anatomy (and its association with LMA), aquaporin activity and enzymatic processes (carbonic anhydrase, RuBisCO) have recently been shown to influence *g_m_* (Muir et al. 2014, Flexas et al. 2006).

In crop plants, the natural abundance of stable N isotopes (δ^15^N) appears to be influenced by soil water availability and together Δ^13^C and δ^15^N, has been proposed as an integrative measure for plant resource use-efficiency (Lopes et al.2006). Lately, there are growing concerns over potential effects of soil-derived inorganic N [ammonium (NH_4_^+^) and nitrate (NO_3_^−^)] affecting riparian systems and Kohl et al. (1971) established a negative relationship between NO_3_^−^ and δ^15^N to provide an insight into N uptake and assimilation by plants. With growing interest to establish *Populus* and *Salix* species as bioenergy crops in riparian buffer systems for nutrient interception and uptake along field edges, it is vital to investigate intraspecific variations in nitrogenisotope discrimination (δ^15^N). Except for the study in *P. balsamifera* by Kalcsits and Guy (2016), no other studies exploited intraspecific variations in δ^15^N. In addition, one could improve nitrogen use efficiency (NUE) by understanding plant N uptake, assimilation and remobilization (between sink and source) during the growing season. These measures of resource use-efficiencies need careful interpretation, as they are confounded to common garden artefacts (Soolanayakanahally et al. 2015).

Seasonal phenologies among deciduous trees in boreal and temperate regions are conditioned by the environment, especially by photoperiod and temperature. Bud phenology of boreal trees is characterized by spring bud break (bud flush) and summer growth cessation followed by leaf senescence in autumn (Fig. 1). The latter two events are cued by photoperiod (Fracheboud et al. 2009) and have an adaptive significance displaying highest heritability (Alberto et al. 2013). As photoperiod regime is precisely the same from year-to-year, one can calculate critical photoperiod from observational data given the calendar date and latitude (Withrow, 1959). For example, Howe et al. (1995) reported that a northern *P. trichocarpa* ecotype (53°N) ceased height growth and set terminal bud to a critical photoperiod of 15h, whereas for a southern ecotype (40°N) the critical photoperiod was 9h.Quite similarly, cold hardiness development displays latitudinal clines during spring and fall in *Tamarix* and *Populus* spp. (Friedman et al. 2011). An earlier bud set among high latitude trees might result in severe infestation of *Melampsora* leaf rust in a common garden setting due to natural selection trade-offs between growth phenology and disease resistance (McKown et al. 2014b).

**Figure 1A.**
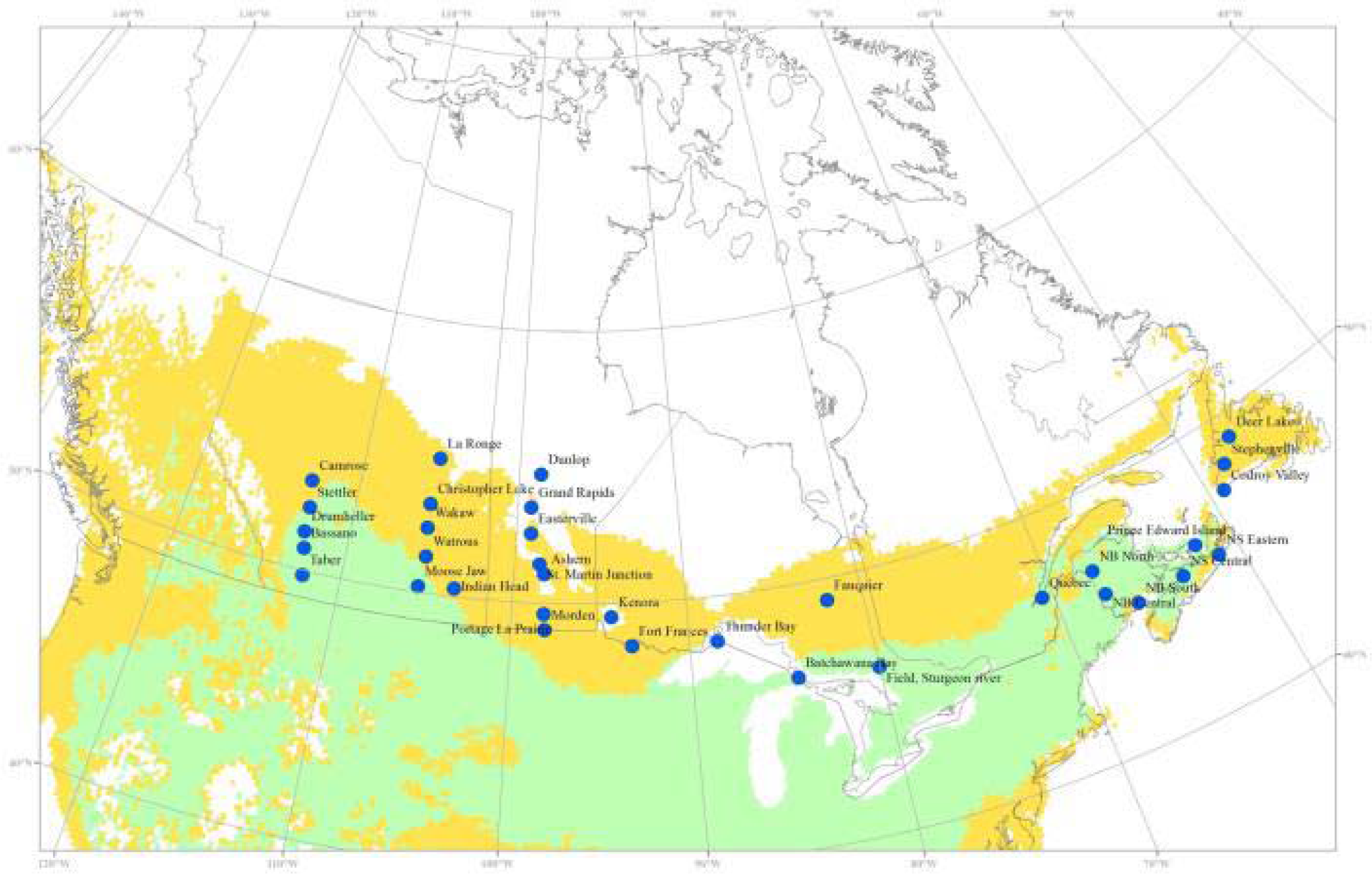
Geographical distribution of 34 native populations (blue circle) of Salix eriocephala from their natural ranges across eastern and western Canada. The green color depicts species dominant continuous range while the yellow shaded area is species discontinues range. The common garden was established at Indian Head, Saskatchewan, Canada.

**Figure 1B.**
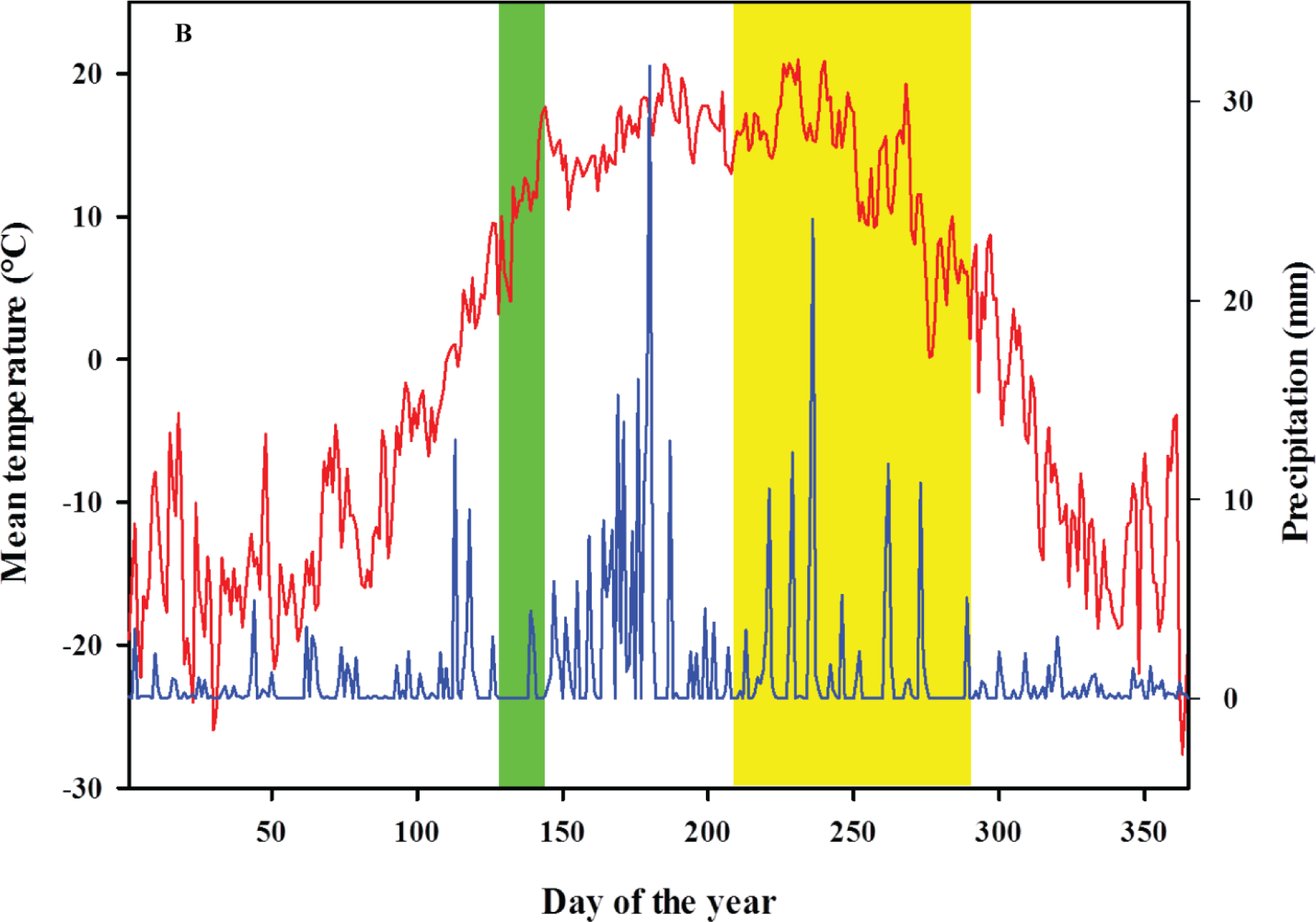
Mean temperature (red line) and precipitation (blue line) that prevailed at Indian Head common garden during 2013 and 2014. The green band represents the spring bud flush period, whereas, the yellow band represents leaf senescence duration among Salix eriocephala populations.

Given the emergence of willows for biomass, bioenergy and environmental applications, many genetic resources are currently being made available in North America (Smart and Cameron 2008) and Europe (Lindegaard and Barker 1997) for detailed investigation of phenotypic variation. One such Canadian genetic resource is the AgCan*Salix* (<u>Ag</u>riculture <u>Can</u>ada <u>*Salix*</u>) collection comprising both native and hybrid willows. Canada has 76 native willow species which are adapted to a wide range of environmental conditions (Argus 2010). The diamond or heart-leaf willow, *S. eriocephala* Michx., spans a broad geographic range couple with diverse climatic conditions (Dorn 1970), whereby selective pressure on growth physiology and seasonal phenology traits is expected to vary extensively.

Considerable intraspecific variation in growth phenology and genetic diversity was documented by earlier studies in *Salix* species that are highly correlated with their latitude of origin and/or growing season length (Weih et al. 2011, Trybush et al. 2012, Berlin et al. 2014, and Pucholt et al. 2015). Quantitative Trait Loci (QTLs) associated with growth phenology traits such as bud burst, elongation growth and leaf abscission were identified in *Salix* spp. (Ghelardini et al. 2014). Through association mapping analysis significant associations for bud burst, leaf senescence and biomass traits were reported in *S. viminalis* (Hallingbäck et al. 2015).

In the present study, we investigated the factors that govern local adaptation by making use of a subset of 34 natural populations of *S. eriocephala.* We hypothesize that the gradient in climatic conditions (frost free days, mean annual precipitation, mean summer temperature) provide selection to favor the populations to respond differentially in functional traits (phenology, photosynthesis, resource use-efficiency) resulting from local genetic adaptation. The specific questions addressed are:

i. Do latitudinal clines exist in photosynthetic assimilation rate (*A*) among genotypes adapted to varying growing season lengths? If so, to what extent stomatal attributes (density and pore length) and mesophyll conductance (*g_m_*) contributes to higher *A*?
ii. Is there variation in water use-efficiency as shown by δ^13^C and nitrogen discrimination as shown by δ^15^N?
iii. Is there a relationship in seasonal phenology and rust infestation among *S. eriocephala* Populations?

## MATERIALS AND METHODS

### AgCan*Salix* collection

During the winter of 2012 (January to April) dormant stem cuttings of *S. eriocephala* were collected from within the natural range of the species within Canada. In total, we sampled 34 populations with 15 genotypes per population (*N*=510, Fig. 1). Care was taken to avoid clonal sampling by selecting distinct genotypes (stools) that were separated by a minimum of 1.0 to 3.0 km apart without phenotypic bias. Dormant cuttings (∼20 cm long) were bagged separately for each of the 34 populations in re-sealable Ziploc^®^ bags and stored at-4°C. Global Information System (GIS) coordinates and other site information were noted foreach genotype (Table 1). This *in situ* collection of *S.eriocephala* along with other native species such as *S. amygdaloides* Andersson., *S. bebbiana* Sarg., *S. discolor* Muhl., *S. interior* Rowlee., and *S. petiolaris* Sm. is commonly referred to as the AgCan*Salix* collection (<u>Ag</u> riculture <u>Can</u>ada *<u>Salix</u>*).

**Table 1.**
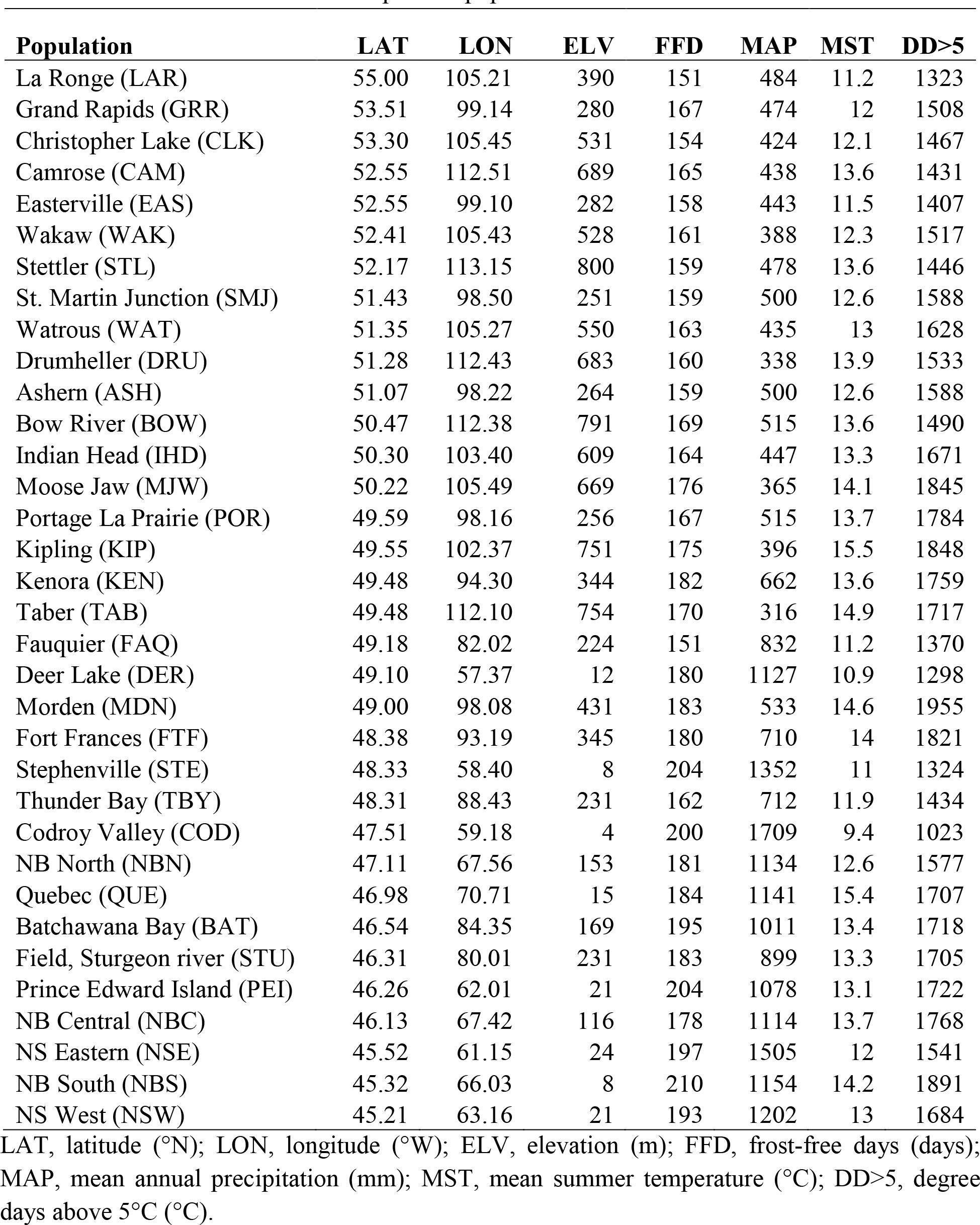
Geo-climatic information of 34 native *Salix eriocephala* populations used in study. The three letters within the brackets correspond to population code.

In spring (mid-May), 8-10 cm long dormant cuttings were forced to root in Spencer-Lemaire rootrainers (Beaver Plastics, Acheson, Canada) using a mixture of Sunshine No.2 (Sun Gro Horticulture, Vancouver, Canada) growing mix (60%), peat (30%) and vermiculite (10%) inside a greenhouse under natural light. Greenhouse conditions were set to day/night temperatures of 23/18 °C, respectively, with relative humidity at40%. The Agriculture and Agri-Food Canada (AAFC) greenhouse facility is located at Indian Head, Saskatchewan (50.52°N 103.68°W;elevation 605 m). Upon bud flush, the plants were regularly watered and fertilized using Hoagland’s solutions at a pH adjusted to 5.8-6.3. After two months of greenhouse growth (∼25-30 cm tall), the plants were transferred to a shade house and allowed to undergo natural senescence.In late-October, the frozen root plugs were individually bagged and stored at −4°C till the following spring.

### Common garden establishment

At Indian Head, the site assigned for the establishment of common garden was left fallow during the 2012 summer. In September, nine soil cores were randomly taken from 0-15 cm depth using an auger to acquire a representative sample along the length of the common garden (3 acres). Upon air drying the soil cores were processed separately, bagged and sent for soil testing at AGVISE laboratories (Northwood, ND, USA). The soil texture was sandy clay loam with an average pH of 7.9 and with 13.6, 19.7, 242.9 ppm of N, Pand K nutrients, respectively. A detailed soil test report is provided in Supplementary table S1.

In spring 2013, the site preparation involved cultivating to a depth ofd20 cm or more and disking. Later, the rows were marked at 3 metre intervals in East-West orientation, with each row running 320 m long. Based on the soil test report, necessary soil amendments were made by drenching the marked rows using micronutrient solutions followed by roto-tilling. Rows were mulched using black plastic sheet (Crawling Valley Plastics, Bassano, AB, Canada) to avoid intra-row weed competition. The frozen root plugs were taken out of cold storage and all fifteen genotypes from each population were planted on mulched rows as a block with 1.0 m spacing. In each of the three replicates, population blocks were randomized with 3 ramets per genotype planted side-by-side. In addition, the common garden hosted a gene bank with a single ramet from 34 populations, totalling to 10 ramets for each genotype (*N*=5100). The site was sprinkler irrigated as necessary during summer months with mechanical weed control between rows. All trees survived the first field growing season. A picture narrative of site preparation and planting is provided in Supplementary Fig. SF1.

### Seasonal growth phenology

Adopting the phenology timetable developed by Saska and Kuzovkina (2010) for *Salix*, bud flush and leaf emergence were monitored in 2014, while leaf senescence and leaf drop were monitored in 2013 and 2014.Phenology was monitored by the same personnel walking through the gene bank every day during spring and twice a week during summer and fall. The *Melampsora* rust disease onset and severity of infection was scored on all genotypes beginning mid-August to late-September following the narrative by Mckown et al. (2014b). Here, we report the *Melampsora* rust scores for the week prior to September 21^st^. Green cover duration (GCD; days) was calculated as the difference between days to leaf senescence and days to leaf emergence. Final height and non-coppiced stem dry biomass were recorded in November 2014. The remaining three replicates were coppiced in November 2013 after initial establishment over first growing season. In the following years (2014 and 2015), phenology was monitored across all three replicates and used for estimating broad-sense heritability (*H*^2^) along with biomass and height gain.

### Screening for morpho-physiological traits

Towards gas exchange measurements, we used a subset of 10 randomly chosen genotypes per population with the exception of Easterville (EAS) where only eight were used (*N*=338). The measurements were made between 5^th^ July and 31^st^ July 2014 during active growth without water deficit as our common garden was installed with a sprinkler irrigation system. All gas exchange measurements were done on clear, sunny days. Briefly, a Li-COR 6400 XTR (LI-COR Biosciences, Lincoln, NE, USA) portable infra-red gas exchange system was used for gas exchange measurements. The gas exchange equipment was switched on by 8:00 am every morning at the common garden location and allowed to stabilize for 30 min prior to recording. On any given day, gas exchange measurements were recorded on a single leaf per genotype between 8:30 and 11:45 am, with the measurement plant randomized among populations and days of measurement. Inside the leaf chamber, the following conditions were maintained: reference CO_2_ concentration set to 400 ppm using CO_2_ cartridges; flow rate 500 μmol s^−1^; block temperature set at 23°C; relative humidity of incoming air adjusted to ˜50-55%; photosynthetic active radiation (PAR) 1000 μmol m^−2^ s^−1^. Maximum photosynthetic assimilation rate (*A*, μmol CO_2_ m^−2^ s^−1^), stomatal conductance (*g_s_*, mol H_2_O m^−2^ s^−1^), transpiration rate (*E*, mmol H_2_O m^−2^ s^−1^) and intercellular CO_2_ concentration (*C_i_*, ppm) were measured on fully expanded leaves. The intrinsic water-use efficiency (*WUE*_i_) was determined by calculating *A/g*_s_ (μmol CO_2_/mmol H_2_O). Later, *A-C_i_* curves were constructed on selected populations representing East (NBN, NBS) and West (CLK, MJW, WAK) using the methodology described by Soolanayakanahally et al. (2009). The maximum carboxylation rate allowed by Rubisco (*V*_cmax_), rate of photosynthetic electron transport (*J*), triose phosphate utilization (TPU) and internal conductance (*g*_m_, mol CO_2_ m^−2^ s^−1^) were estimated by fitting the *A-C_i_* curve data to the model of Sharkey et al. (2007).

Following gas exchange measurements, chlorophyll content index (CCI) was measured on three fully expanded leaves per plant using an Opti-Sciences CCM-200 meter (Hudson, NH, USA) and averaged for statistical analyses. Two leaf discs were sampled using a hand held paper punch exactly from the same leaf used for gas exchange, and oven dried at 50°C for 72 h for recording leaf mass per unit area (LMA, mg mm^−2^). The stem wood was collected in November 2014 at 15cm above ground after recording non-coppiced biomass. Later, these leaf discs and stem wood samples were used to analyze leaf and wood carbon (C) and nitrogen (N) content and stable isotopes ratios (δ^13^C and δ^15^N; %o) at the UC Davis Stable Isotope Facility (Davis, CA, USA). All δ^13^C values were converted to Δ^13^C using Farquhar et al. (1989) equation with isotopic composition of the air to PeeDee Belemnite of −8.3‰. Leaf C to N ratio (C: N; mg/mg) and photosynthetic nitrogen-use efficiency (PNUE; prnol CO_2_ g^−1^N s^−1^) were calculated from these values. Stomatal density (number of stomata per unit of leaf area, mm^−2^) measurement samples were prepared by applying a thin coat of clear nail polish on the adaxial and abaxial surfaces of fully expanded leaves (Gornall and Guy 2007). The dried impressions were stripped from leaves and mounted onto clear microscopic slides for observation. The slides were viewed under Zeiss phase contrast microscope (Axio Lab A.1, Toronto, ON, Canada) and stomata were counted under 20× magnification. The final stomatal count was averaged from three randomly selected field views from one impression. The stomatal pore length (μ,m) was measured on a subset of populations based on density ranks representing East (high stomatal density: PEI, NSW, QUE, NBS) and West (low stomatal density: MDN, DRU, KEN, IHD). Five genotypes from each population (*N*=40) were randomly chosen to determine pore length on 5 stoma per genotype (*N*=200). Later, maximum stomatal conductance to CO_2_ (*g*_c(max)_, mol m^−2^ s^−1^) was estimated using a modified version of the Franks et al. (2012) equation.

The Climate Normals (1981-2010) of closest stations were obtained for all populations from Environment Canada (http://www.climate.weatheroffice.ec.gc.ca/climatenormals/indexe.html). Climate variables included frost free days (FFDs, days), mean annual precipitation (MAP, mm), mean summer temperature beginning from May to September (MST, °C) and degree days above 5°C (DD>5, °C). The FFD was calculated based on the number of days where minimum day temperature was above 0°C, a proxy for growing season length at each location (Table 1).

### Statistical analysis

All statistical analyses were performed using *R* studio statistical software (0.99.484 for Windows). Wherever possible the data from growth phenology traits were calculated from pooled data across 2013 and 2014. Analysis of variance (ANOVA) and correlation analysis for traits were performed to estimate the functional trait diversity and relationship among the populations used in the study. Broad-sense heritability (*H*^2^) estimates of traits were calculated as a ratio ofgenetic variability (σ^2^ _g_) to phenotypic variability 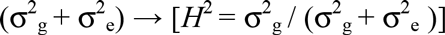 Variance components were estimated based on ANOVA, where σ^2^ _g_=M_e_ and 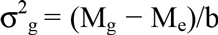 [M_e_; mean square of error, M_g_; mean square of genotypes and b; number of replications].

Pearson’s correlation was performed to estimate correlation coefficients (*r*) on all 338 genotypes among morpho-physiological traits and geo-climatic variables. Significant correlations between traits were expressed after Bonferroni correction (*P*<0.001).

## RESULTS

The major aim of this study was to evaluate the extent of intraspecific variation in growth physiology and seasonal phenology of *S.eriocephala*; hence, genotypes were selected to be representative of a broad range of latitudes and longitudes. The geographical range spanned 15° in latitude and 52° in longitude, with elevation ranging from 4 to 800 m (Table 1). In general, the species range for *S.eriocephala* is at higher latitudes in the west, causing similar associations with longitude and elevation across Canada (Fig. 1). The average number of frost free days (a proxy for growing season length) ranged from 151 to 210 days with precipitation increasing from West (316 mm) to East (1709 mm). Growing degree days (DD>5 °C) were 60 units higher for eastern genotypes. A total of 25 traits related to ecophysiology, phenology and biomass were measured in 338 genotypes sourced from 34 populations.

### Phenotypic trait variations in *S.eriocephala*

All measured traits showed a wide range of variation between genotypes and among populations (Table 2). Photosynthetic assimilation rates (*A*) ranged from 9.1 to 23.7 μmol CO_2_ m^−2^ s^−1^, with intrinsic water-use efficiency (*WUE*_i_) fluctuating between 19.5-128.7 prnol CO_2_/mmol H_2_O. Both LMA and stomatal density displayed large variations among genotypes. Overall, leaf Nand leaf C:N ratios were higher than wood, with CCI ranging from 6.5 to 22.2 units. The carbon isotopic discrimination (Δ^13^C) for leaf and wood ranged from 16.8 to 23.1‰. While, the nitrogen isotopic discrimination (δ^15^N) ranged between 3.4 to 21.2‰ in leaf and wood tissues. Spring bud flush occurred within a week’s time, whereas, leaf senescence spanned over 80 days. At our common garden location, the green cover duration ranged from 98 days to 166 days leading to variances in overall height gain and biomass accrual.

**Table 2.**
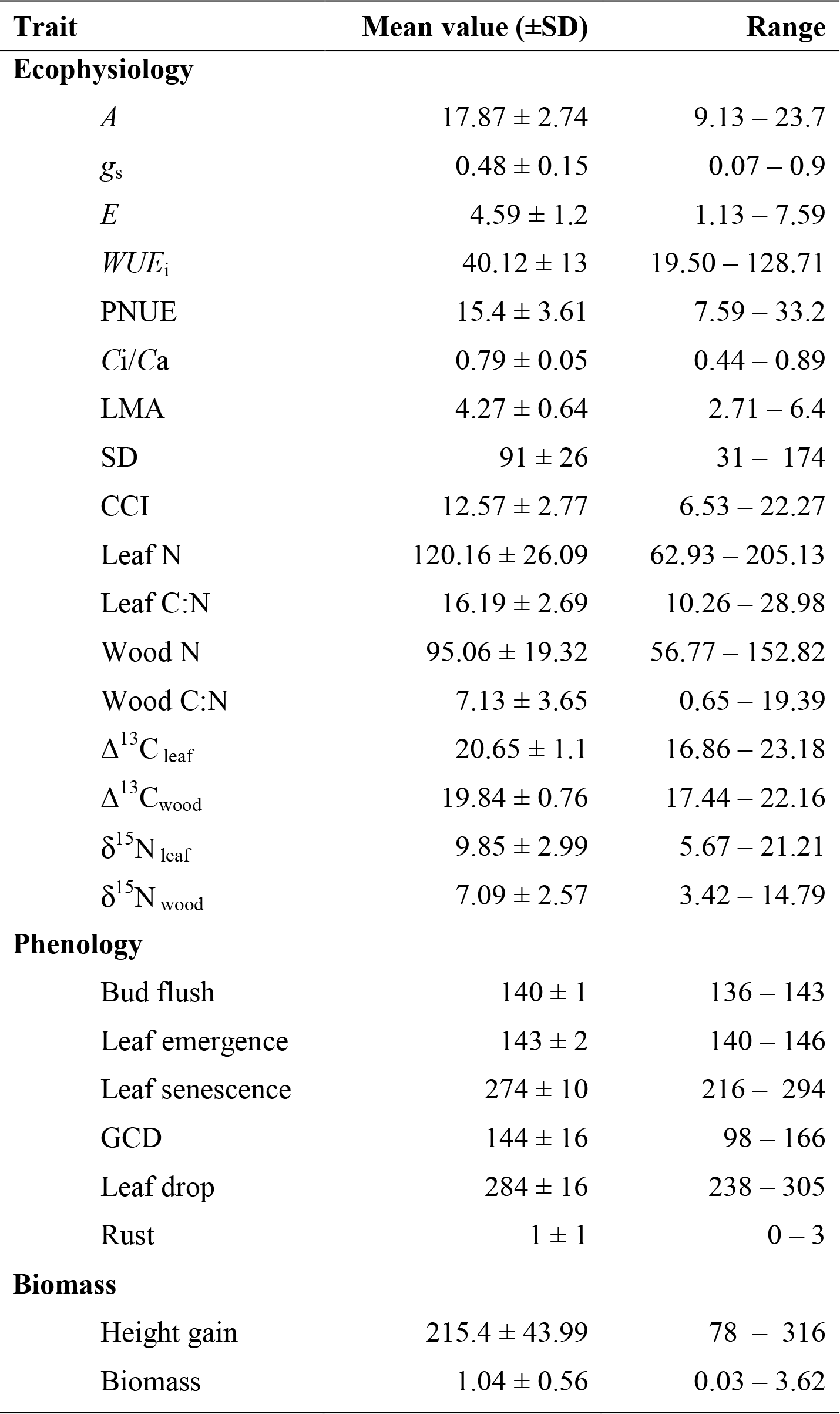
Mean values for ecophysiology, phenology and biomass traits measured in 338 genotypes of *Salix eriocephala.*

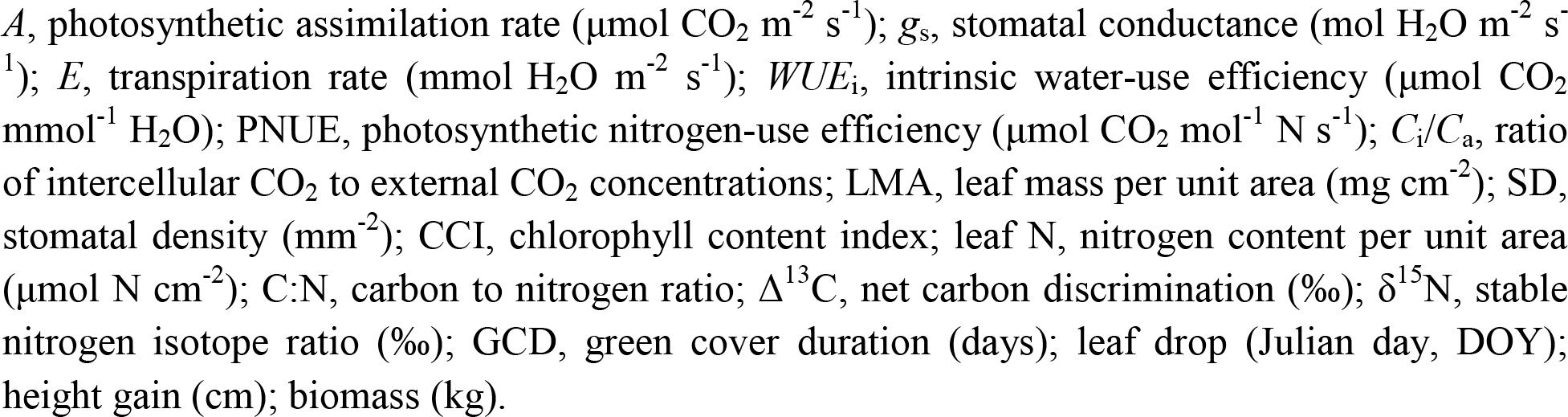

### Correlation between geo-climatic and phenotypic variables

Pearson correlation coefficients (r) between geo-climatic parameters and phenotypic traits for all 338 *S.eriocephala* genotypes are shown in Table 3. It is evident that both *A* and g_s_ increased with increase in latitude (LAT), longitude (LON), elevation (ELV) and MSP. Conversely, both traits were negatively correlated with FFD and MAP. *WUE*_i_ among genotypes increased with increase in FFD and MAP. LMA increased with increase in LAT and LON implying that the leaves got thicker at high latitudes. While, stomatal density followed opposite trend. Both CCI and leaf N increased with increase in LAT, LON and ELV and decreased with increase in FFD and MAP. δ^15^N_leaf_ and 5^15^N_wood_ paralleled above trends. PNUE, wood N, Δ^13^C_leaf_ and Δ^13^C_wood_ were not significantly affected by any geo-climatic variables. The relationships between LAT and resource use-efficiencies [water (Δ^13^C) and nitrogen (δ^15^N)] are plotted in Fig. 4A and 4B, respectively. The δ^15^N values of both leaf and wood increased significantly with increase in LAT (Fig. 4B). The DD>5°C did not have any significant influence either on gas exchange orphenology traits.

**Figure 2.**
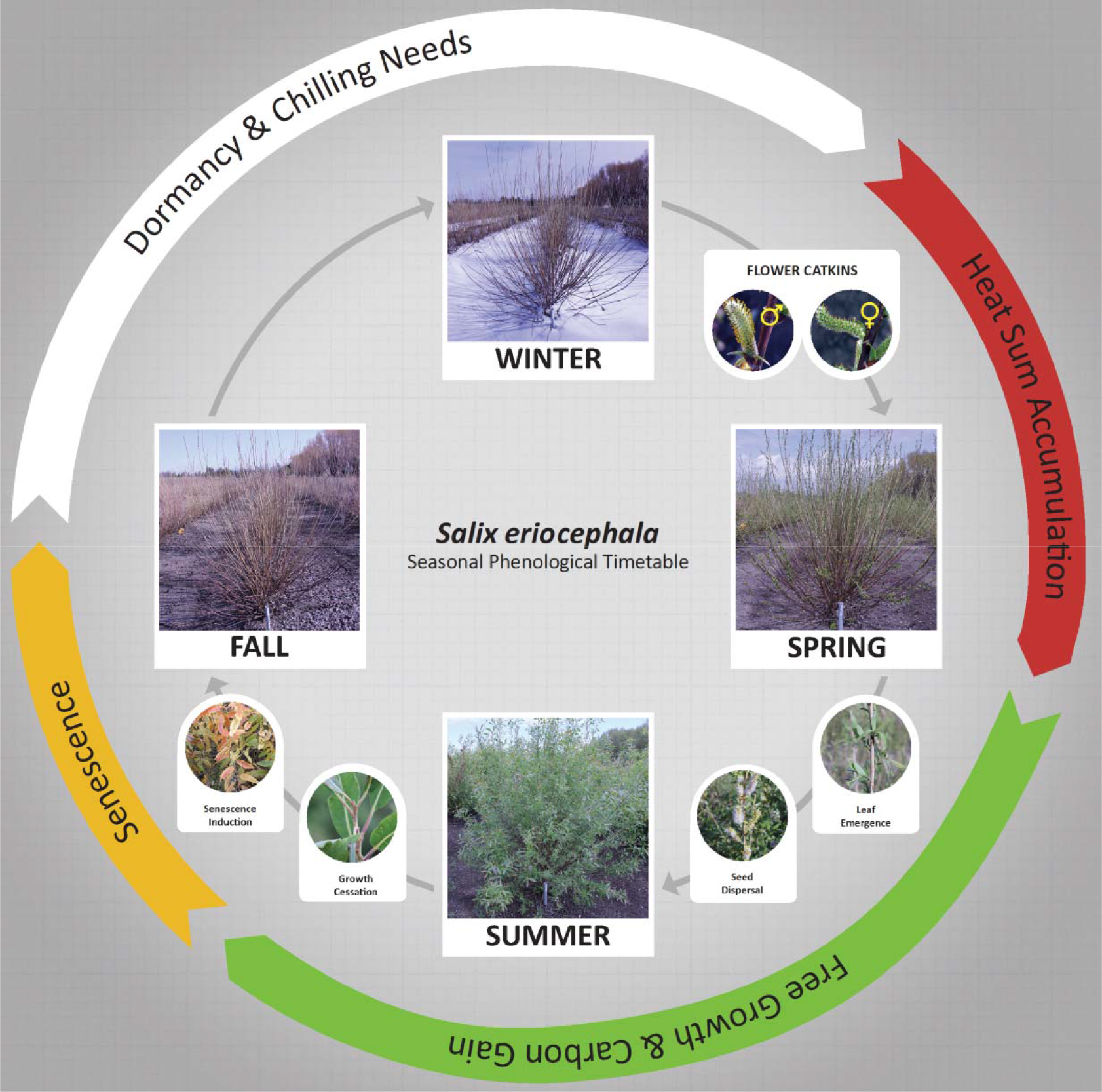
Seasonal phenological timetable of *Salix eriocephala*. Plants become dormant in autumn (fall) and remain so until chillingrequirements are during winter (white band); during spring, heat sum accumulation leads to bud flush and leaf emergence (red band); free growth and carbon gain occur after leaf emergence (green band); growth cessation and leaf senescence during late summer and early autumn (yellow band).

**Figure 3.**
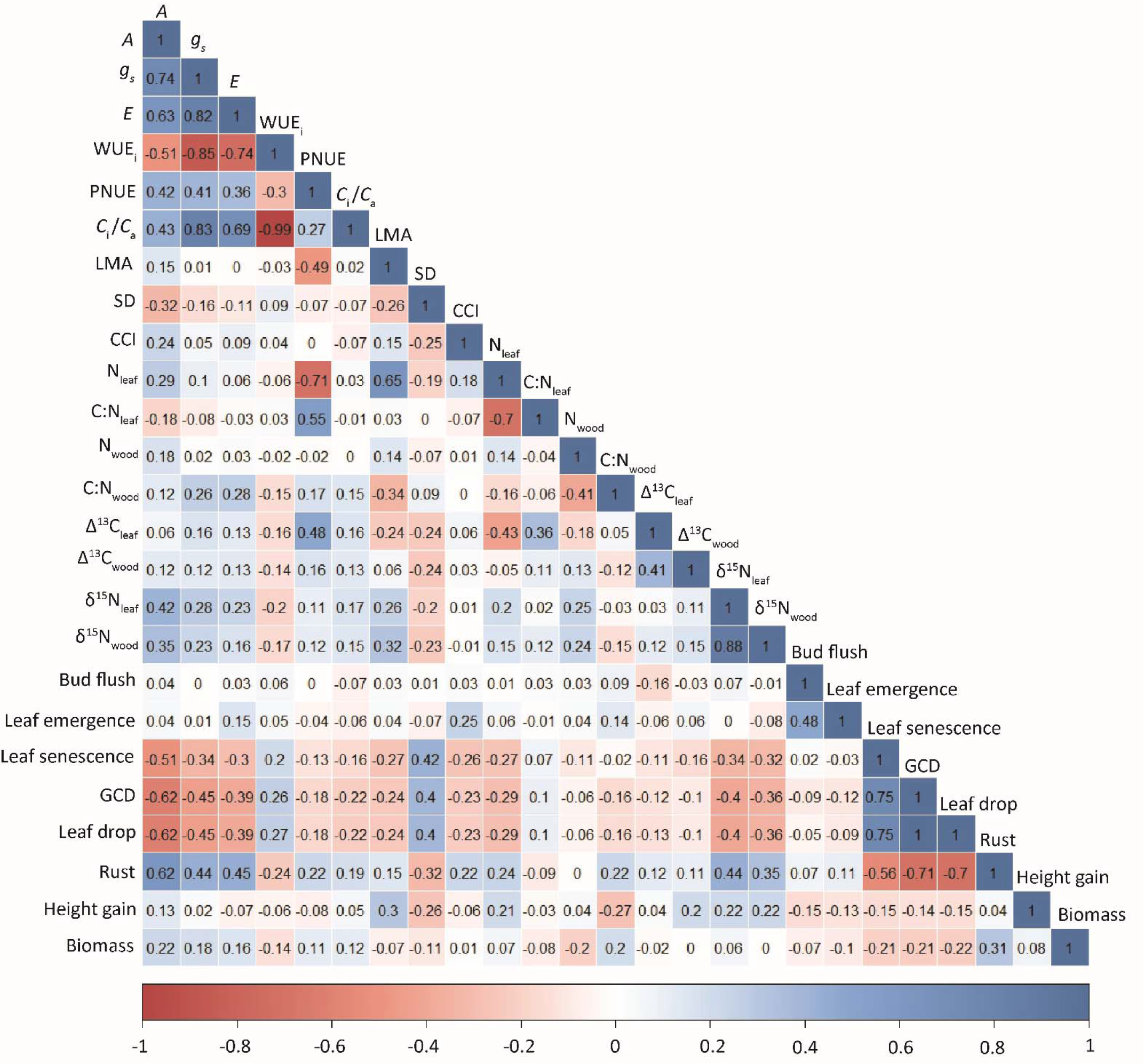
Heat map of Pearson’s correlation coefficient for phenotypic traitsamong 338 populations of *Salix eriocephala*. The scale barbeneath the heat map denotes the direction/magnitude of correlation between the traits, 1 indicated by dark blue being positive and-1 indicated by dark red being negative.

**Figure 4.**
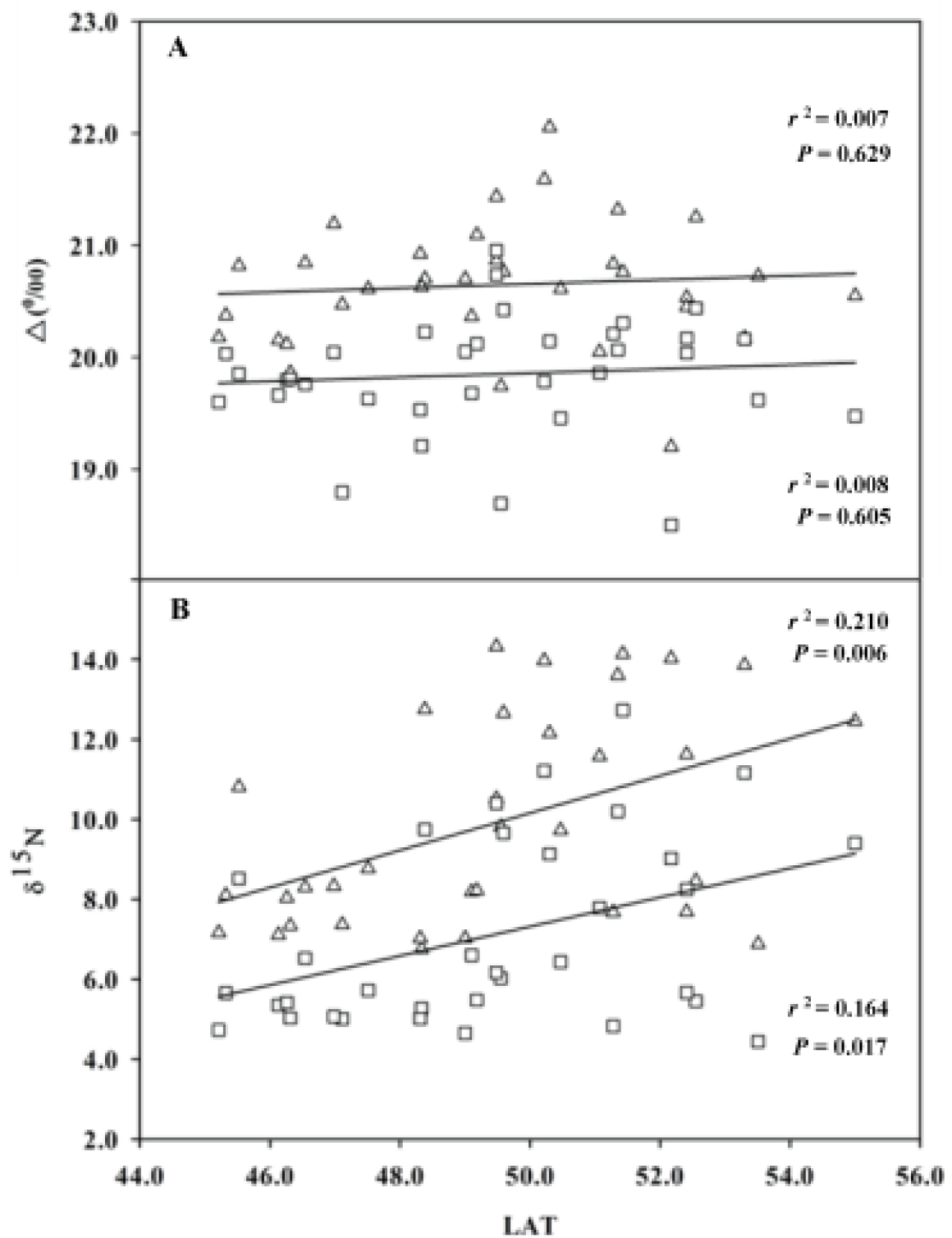
**A**.Carbon isotope discrimination as determined on leaves (Δ^13^C_leaf_) (Δ) and wood (Δ^13^C_wood_) (Ↄ) and **B**. Nitrogen isotope composition of leaves (δ^15^N_leaf_) (Δ) and wood (δ^15^N_wood_) (Ↄ) of *Salix eriocephala* populations plotted against their latitude of origin.

Spring phenology (bud flush and leaf emergence) did not correlate with any geo-climatic variables. However, autumn events (leaf senescence, leaf drop, GCD) were negatively correlated with LAT, LON and ELV and positively correlated with FFD and MAP. The *Melampsora* rust incidence among the populations followed the reverse trend. Rust onset and its severity were positively correlated with LAT, LON, ELV and MST, but were negatively correlated with FFD and MAP. Genotypes from higher latitudes made significant height gains displaying positive correlations with LAT and LON.Non-coppiced single stem biomass was negatively correlated with FFD and MAP in our common garden.

### Correlation among phenotypic variables

Traits related to ecophysiology, phenology and biomass were analyzed for functional intercorrelations and shown in a heat map (Fig. 3). The ecophysiological traits analyzed among the populations showed significantly stronger positive or negative correlations among each other. Photosynthetic assimilation rate (*A*), stomatal conductance (*g*_s_), transpiration rate (*E*), PNUE and *C*_i_/*C*_a_ were largely positive correlated to each other. The *WUE*_i_ was negatively correlated with *A*, *g*_s_, *E* and *C*_i_/*C*_a_. LMA is negatively correlated with PNUE and positively with Leaf N, δ^15^N_leaf_, δ^15^N_wood_ and height. Stomatal density was positively correlated with autumn phenology and negatively with *A*, rust and height gain. CCI and leaf N are positively correlated with *A* and rust. Δ^13^C_leaf_was correlated positively with PNUE and negatively with leaf N. δ^15^N_leaf_ and δ^15^N_wood_ were positively correlated to each other and also with *A*.

Most of the phenological traits were negatively correlated to ecophysiological traits (Fig.3). Bud flush was not significantly correlated to either *g*_s_ or PNUE. A similar trend was observed between leaf unfolding and δ5^15^N_leaf_ and between rust incidences. Leaf senescence,GCD and leaf drop were negatively correlated to *A*, *g*_s_, *E*, CCI, leaf N, δ^15^N_leaf_, δ^15^N_wood_ and rust incidence, and positively correlated to *WUE*_i_ and stomatal density. The correlation between leaf senescence, GCD and leaf drop was significantly positive (*r*=0.75). Height and biomass showed less correlation with ecophysiological and phenology traits. There was no significant correlation between biomass and Δ^13^C_wood_ and δ^15^N_wood_.

### Stomatal pore length and *g*_m_

Based on the stomatal density ranking we measured stomatal pore length in eight populations representative of either ends of the range. Populations originating from Eastern Canada (low latitudes) had high stomatal density (124±3.08 SE) per unit leaf area with a smaller pore length (11.168 μm). In contrast, populations originating from Western Canada (high latitudes) had fewer stomata (68±2.19 SE) per unit leaf area but a longer pore length (16.864μm). The significant differences in stomatal pore length and stomatal density are shown in Fig. 5A-5B (*P*<0.001). We also observed significant differences in *g*_s_ (Fig. 5C, *P*<0. 001) among eight populations, however, the maximum diffusive conductance to CO_2_ (*g*_c(max)_) as determined by stomatal density and pore length reached physiological optima at either ends ofthe species range to achieve maximum carbon gain (Fig. 5D, *P*=0.09).

**Figure 5.**
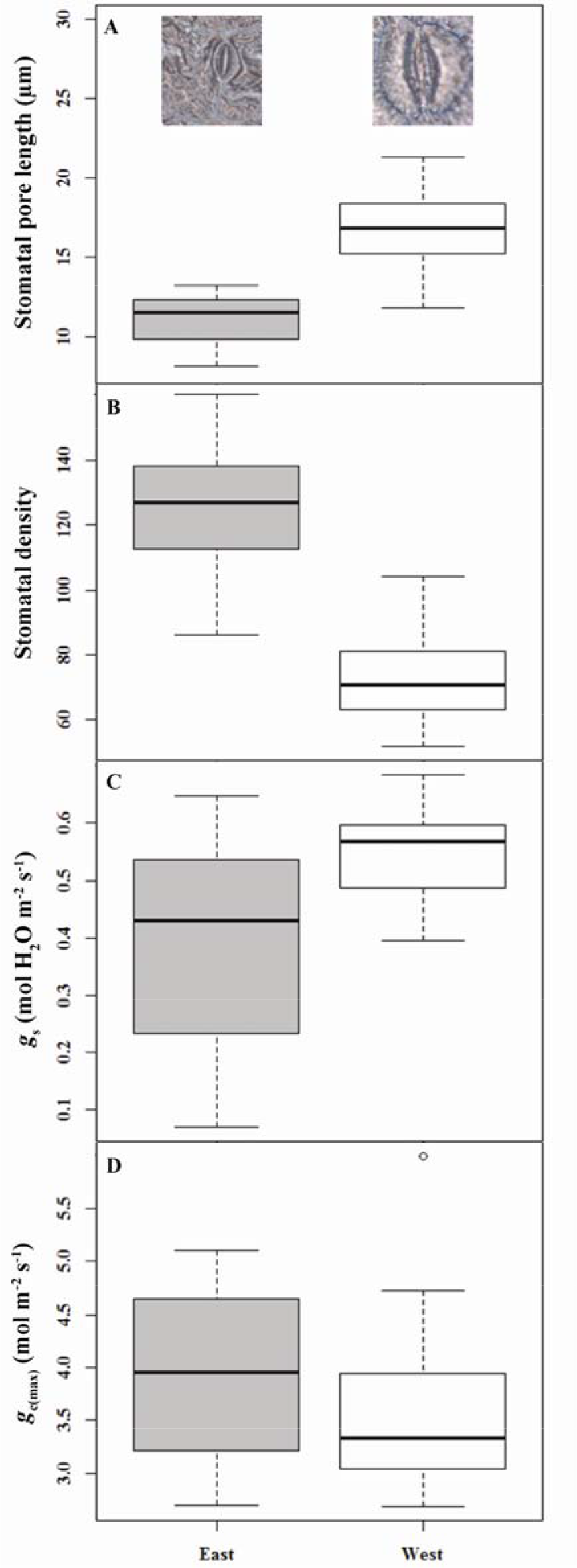
The relationship between stomatal dimensions and maximum leaf condutcances of selected eastern and western populations of *Salix eriocephala*. **A**. Mean stomatal pore length (SL,μm, images above the boxes represent stomatal pore length from respective latitudes viewed under 100×magnification using phase contrast microscopy), **B**. mean stomatal density (SD), C. stomatal conductance to carbon dioxide and water vapor (*g*_s_, mol H_2_O m^−2^ s^−1^) and D. maximum diffusive conductance to carbon dioxide (*g*_c(max)_, mol m^−2^ s^−1^) as determined by stomatal dimensions of eastern (NBS, NSW, PEI and QUE) and western (DRU, IHD, KEN and MDN) populations.

Following gas exchange measurements, a total of ten CO2 response curves were constructed and data analyzed using *A*-*C*_i_ curve fitting model. The corresponding estimates of *g*_m_, *V*_cmax_, *J* and TPU were plotted in Fig. 6A-6D for three populations from the west (CLK, MJW, WAK) and two populations from the east (NBN, NBS). The western genotypes had higher g_m_ (0.288 *vs*. 0.198; mol CO_2_ m^−2^ s^−1^), *V*_cmax_ (107 *vs*. 93), *J* (146 *vs*. 114) and TPU (10.52 *vs*. 8.48) values than the eastern genotypes. The variance between their mean values shows the difference in magnitudes of these values.

**Figure 6.**
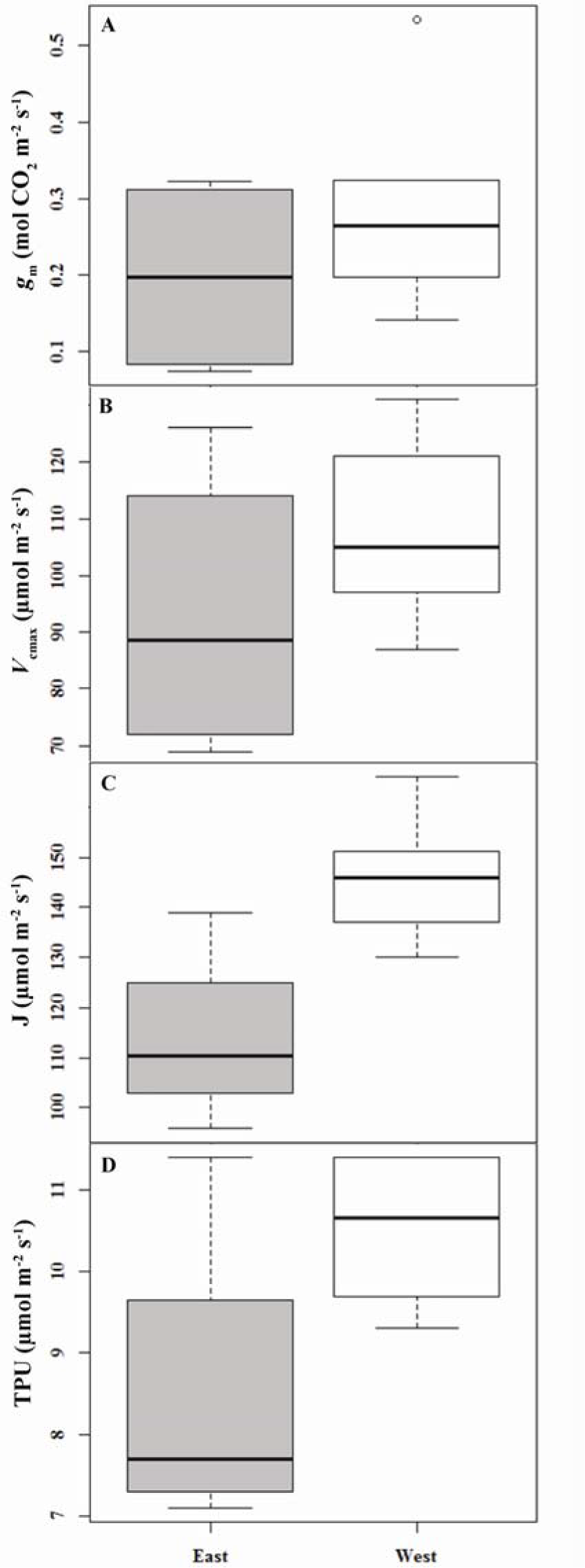
Gas exchange traits of selected eastern and western populations of *Salix eriocephala*. **A**. Internal conductance (*g*_m_, mol CO_2_ m^−2^ s^−1^), **B**. Maximum carboxylation rate allowed by rubisco (*V*_cmax_, μmol m^−2^ s^−1^), C. rate of photosynthetic electron transport based on NADPH requirement (*J*, μmol m^−2^ s^−1^) and D. Triose phosphate use (TPU, μmol m^−2^ s^−1^) of eastern (NBN and NBS) and western (CLK, MJW and WAK) populations estimated using Sharkey’s *A*-*C*_i_ curve fitting model. *J* was significantly different (*P*<0.05) between the eastern and western populations.

### Broad-sense heritability and rust infestation

Heritability estimates (H^2^) estimates were calculated for seasonal phenology and biomass related traits. The *H* estimatesranged from 0.62 to 0.95 among the traits (Table 3). Bud flush, leaf emergence, leaf senescence and leaf drop had heritability estimates of 0.72, 0.77, 0.62 and 0.78, respectively. Height gain had higher heritability estimate (0.95) than biomass (0.88). Alatitudinal cline with *Melampsora* rust incidence scores are shown in Fig. 7.

**Table 3.**
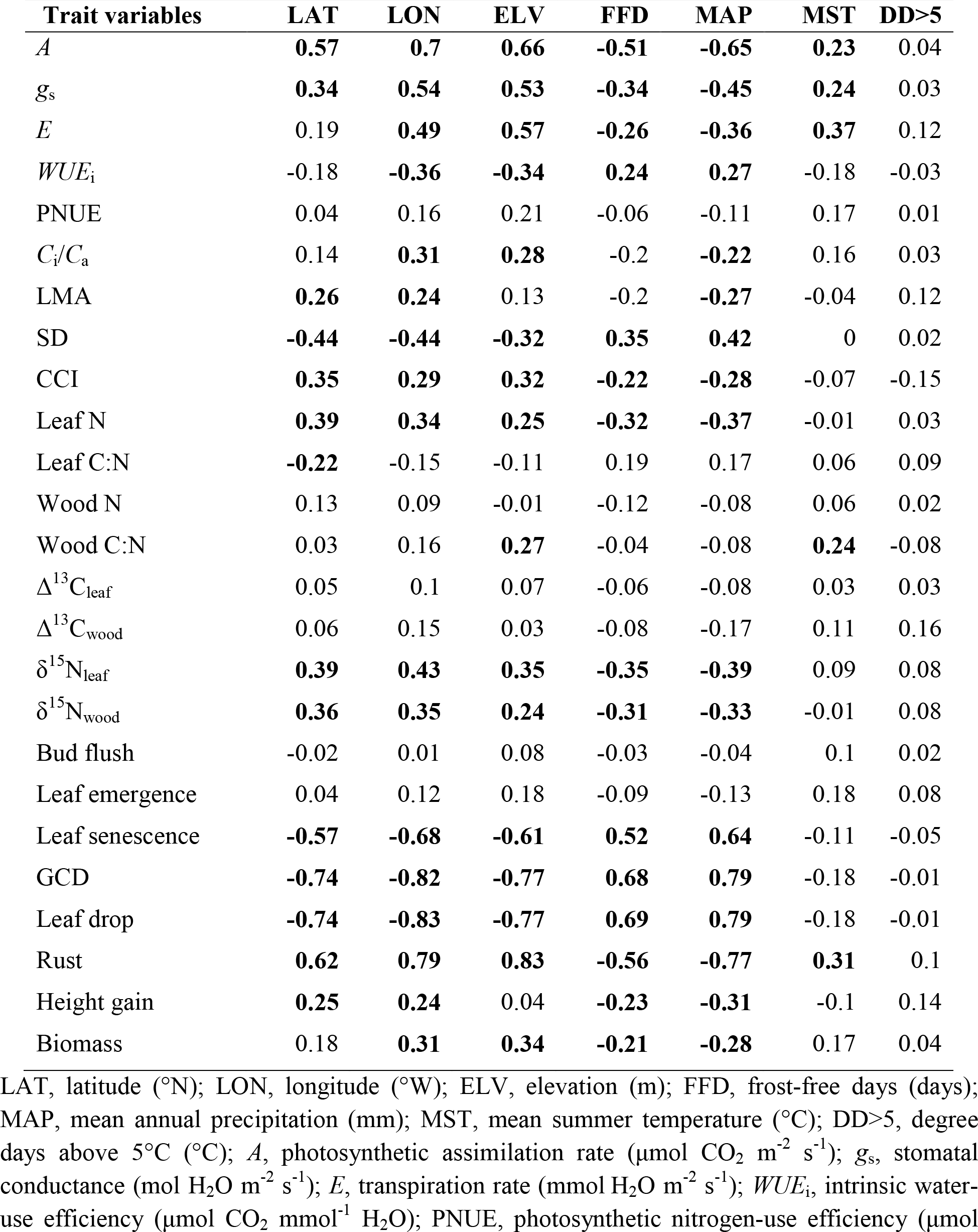
Pearson correlations coefficients (*r*) betweengeography, climate and physiological variables for all 338 genotypes. Boldare significant after Bonferroni correction (*P*<0.001).

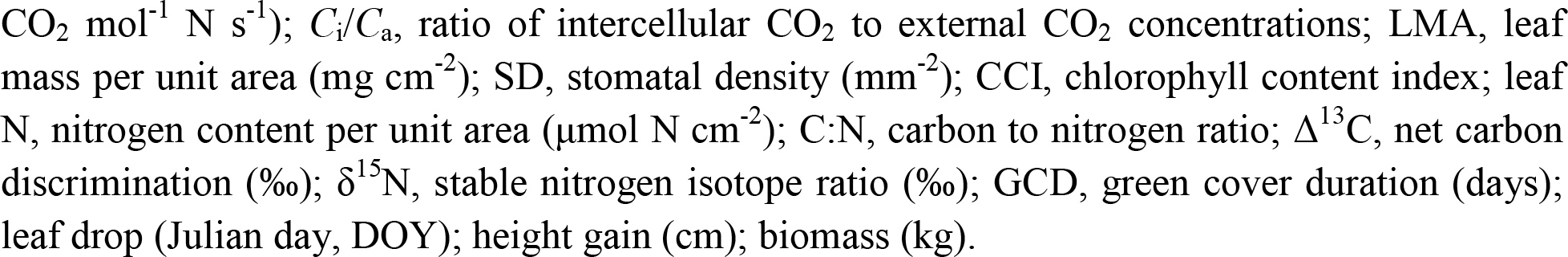

**Figure 7.**
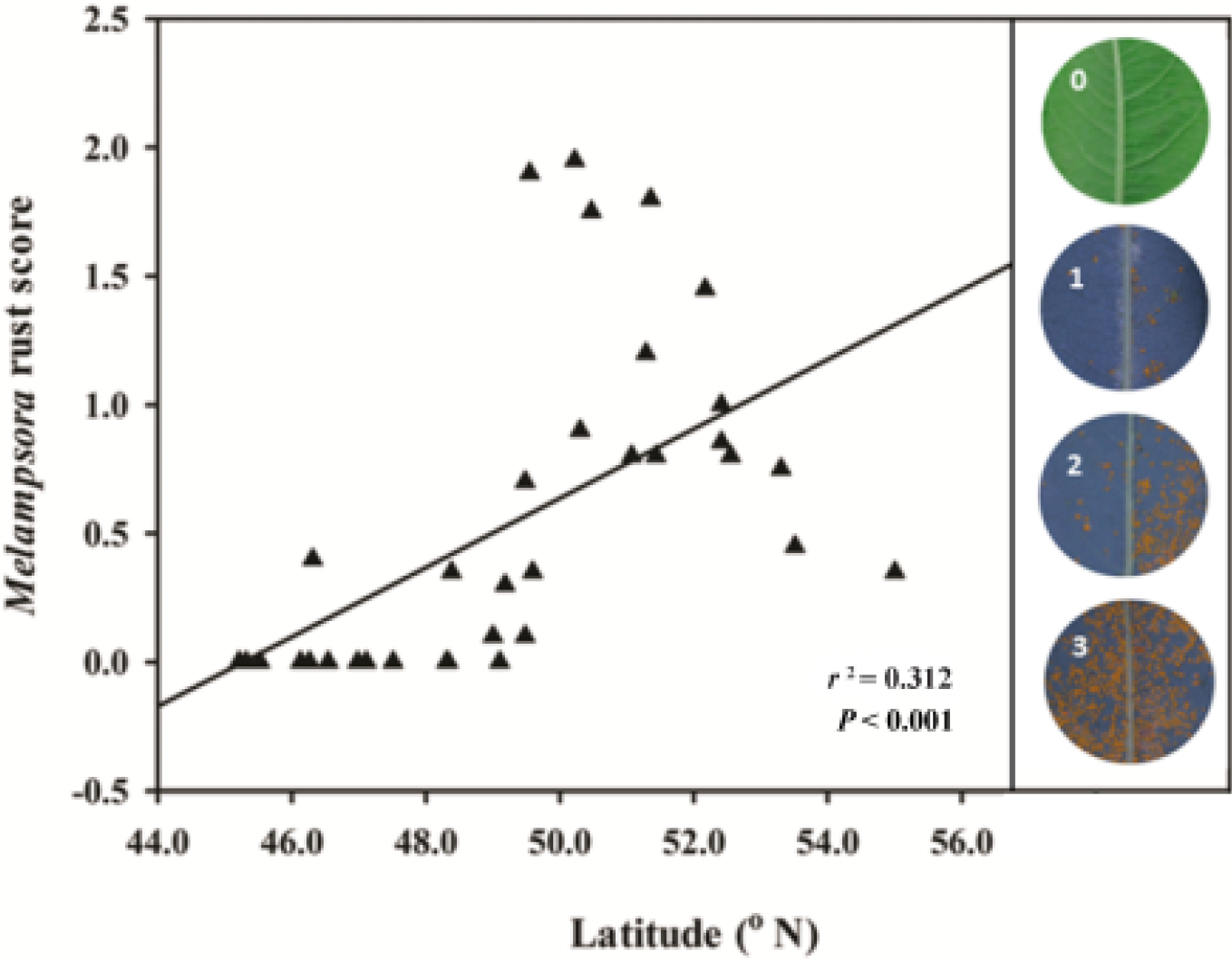
*Melampsora* rust incidence score of *Salix eriocephala* populations plotted against their latitude of origin. The images indicate the rust incidence scoring key, 0 for no rust and 1, 2 and 3 for minimum, moderate and maximum rust symptoms in the leaves,respectively.

**Table 4.** Broad-sense heritability (*H*_2_) estimates of phenology and biomass traits in 2015.

| Traits | Heritability ($H^2$) |
| --- | --- |
| Bud flush | 0.72 |
| Leaf emergence | 0.77 |
| Leaf senescence | 0.62 |
| Leaf drop | 0.78 |
| Height | 0.95 |
| Biomass | 0.88 |

## DISCUSSION

This is the first comprehensive study on a substantial number of native populations that originated from varying latitudes and longitudes in *S. eriocephala*. The above results emphasize the importance of considering key ecophysiological and phenological traits while studying local adaptation among willow populations of divergent origin. Besides, these physiological mechanisms are discussed in the light of convergent evolution among the members of the genus *Salicaceae*-a sympatric adaptive phenotype to very similar climate and photoperiod.

### Adaptive variations in photosynthesis

Planted into a common environment (indoor greenhouse and/or outdoor common garden), observed functional traits differences among populations originating along an environmental gradient can be influenced by past evolutionary history resulting in adaptive genetic variations. In our study, photosynthetic assimilation rate (*A*) increased with increase in latitude when measured during free growth in *S.eriocephala* genotypes. This is an agreement with the previous findings in North American *Populus* species which occupy similar climates (Gornall and Guy 2007, Soolanayakanahally et al. 2009, McKown et al. 2014a, Kaluthota et al. 2015). Collectively, these studies hypothesised that the observed patterns of higher *A* among high latitude genotypes represent true adaptive variation in response to growing season length. Whereby, genotypes from shorter growing seasons possess inherently higher *A* compared to the genotypes from longer growing seasons. Again, higher *A* in *S.eriocephala* is associated with higher *g*_s_, LMA, CCI and Leaf N (Soolanayakanahally et al. 2009, McKown et al. 2014a, Kaluthota et al. 2015), resulting in greater height gain among high latitude genotypes. A similar height gain was observed among high latitude *P.balsamifera* populations when photoperiodic constraints were removed by growing under extended daylength (Soolanayakanahally et al. 2009). On the other hand, when daylength was limiting, height rankings reversed leading to alterations in root:shoot ratios (Soolanayakanahally et al. 2013).

Significant increases in *A* were found in other deciduous tree species (Benowicz et al. 2000, Soolanayakanahally et al. 2015) and evergreen conifers (*Picea glauca* [Moench] Voss, Benomar et al. 2015) sourced along a north-south gradient with varying growing season length. Such adaptation to growing season length in photosynthetic assimilation rates can be generalized along elevational gradients as well (Oleksyn et al.1998).

Plant species occupying large geographic areas provide cues about adaptation mechanisms to various environmental conditions (Brosché et al. 2010). Leaf stomata regulate CO_2_ uptake and H_2_O use during photosynthesis and transpiration, respectively. Our *S.eriocephala* genotypes were entirely hypostomatous and stomatal density is negatively correlated with latitude and longitude (while, *g*_s_ is positively correlated with latitude and longitude). Stomatal density and pore length were negatively correlated with each other, one compensating for the other. Wang et al. (2015) studied latitudinal variation in stomatal traits across 760 species to highlight a strong negative relationship between stomatal density and stomatal length governing physiological adaptation to the environment. In their study, the plant species at low latitudes had higher stomatal density and reduced stomatal length than those distributed at high latitudes.

Among plant groups, the maximum diffusive conductance to CO_2_ (*g*_c(max)_) and water vapor is ultimately determined by stomatal density and pore length which may serve as a physiological framework to optimize leaf carbon/water balance (Franks et al. 2012). These long-term evolutionary scale adjustments in stomatal density and porelength in response to environmentalconditions have facilitated *S.eriocephala* to expand into newer habitats leading to local adaptation. As the epidermal stomatal design features evolved 400-million years before present, the observed negative relationship between stomatal density and pore length suggests a widespread highly conversed genetic basis among vascular plants (Franks and Beerling 2009). So, we postulate that larger pore length with fewer number of stomata per unit leaf area certainly contribute to high *g*_s_, in turn higher *A* among high latitude genotypes-a necessary“*energy* constraint *trade-off*” to maximise returns in *g*_s_ and *A* for a given investment in stomata construction costs under global leaf economics spectrum.

The observed difference in *A* is determined by *g*_s_ resulting from a combination of stomatal density and pore length; however, we must not discount the role of *g*_m_ as well. The physiological mechanisms involved in higher *A* have been the subject of extensive investigation in the recent past (Muir et al. 2013, Cano et al. 2013, Buckley and Warren 2014, Barbour et al. 2015). Such an examination of underlying physiological mechanisms in *S.eriocephala* are none and this study provide a first glimpse into naturally occurring variability in *g*_m_. Even though our results on a small subset were not significant for *A*-*C*_i_ curve fitting estimates, overall the observed trends points towards higher *g*_m_ at high latitudes. Previously, Soolanayakanahally et al. (2009) reported an adaptive role of *g*_m_ in enhancing CO_2_ uptake efficiency and photosynthetic capacity among *P.balsamifera* trees adapted to short growing season. This enhanced *g*_m_ is linked to increased palisade surface area exposed to intercellular air space for CO_2_ diffusion (Milla-Moreno et al. 2016), accounting for the positive association between LMA and *g*_m_ (Ryan 2015). Such positive and negative association between LMA (thicker or denser leaf tissue, or both) and *g*_m_ is reported in other plant species as well. In addition, Théroux Rancourt et al. (2015) highlighted the importance of mesophyll-to-stomatal (*g*_m_/*g*_s_) ratios while breeding for dry climates within *Salicaceae* species. Finally, we provide evidences for higher *A* among high latitude *S.eriocephala* genotypes ably supported by higher *g*_s_, larger stomatal pore length and enhanced *g*_m_.

### Clines in resource acquisition

Establishing the link between resource acquisition efficiencies and ecophysiological traits among the populations is essential to further understand their adaptive behaviours. Díaz et al. (2016) mapped global trait spectrum in 46,085 vascular plant species to reflect“acquisitive *vs*. conservative” trade-offs between LMA and leaf N in constructing photosynthetic leaf. In their study, across biomes and plant species, cheaply constructed leaves with short lifespan were nitrogen-rich with low-LMA (acquisitive leaves), while leaves with long lifespan were nitrogen-poor with high-LMA (conservative leaves). Conversely, we found opposite patterns in LMA within *S.eriocephala* collection that occupies temperate-boreal climates. Whereby, shorter lifespan leaves had higher LMA (nitrogen-rich), while longer lifespan leaves had lower LMA (nitrogen-poor). Similar“with in species”patterns in LMA was also observed in *P. balsamifera* that encompass vast geographic ranges (Soolanayakanahally et al. 2009). Even though no significant association exists between LMA and ELV in our study, others found that LMA increases with ELV as well (Poorter et al. 2009).

Higher leaf N contents are associated with higher *A* as large amounts of inorganic nitrogen (∼75%) are present in the chloroplast (Evans and Seemann 1989). We observed a strong positive correlation between *A*,leaf N and LMA. At the same time, PNUE is negatively associated with leaf N and LMA. So possible explanations for high LMA to have lower PNUE could be due to variation in nitrogen allocation between photosynthetic vs. non-photosynthetic structures, and also as a result of differential allocation of photosynthetic N between light harvesting complexes, electron transport and CO_2_ fixation (Field and Mooney 1986). It seems that low latitude *S.eriocephala* genotypes invest more N towards foliar structures to withstand biotic and abiotic stressors, while fast-growing high latitude genotypes allocates more N to photosynthetic apparatus. Previously, Weih and Rönnberg-Wästjung (2007) concluded a positive association between leaf N and photosynthetic capacity in *Salix* genotypes.

*WUE*_i_ decreases as *C*_i_/*C*_a_ increases, suggesting a potential intrinsic trade-off between *WUE*_i_ and PNUE (Field et al. 1983). Both these resource use-efficiency indicators of gas exchange (WUE_i_ and PNUE) mutually depend on *g*_s_, and are influenced by leaf-to-air temperature, light and available soil moisture. Unlike H_2_O, CO_2_ faces further resistance in diffusion from intercellular spaces to the site of carbon fixation (*g*_m_). But, when *P.balsamifera* was grown without resource limitation, *WUE*_i_ increased with increase in latitude (Soolanayakanahally et al. 2009). We recognise the limitation in inferring *WUE*_i_ based on a single common garden. Turner et al. (2010) were able to differentiate the genetic and plastic responses in Δ^13^C in *Eucalyptus* species by taking into account the results from two common gardens.

A negative relationship between *WUE*_i_ and Δ^13^C has been extensively reported in many plants and is genetically determined (Farquhar et al. 1989). Δ^13^C values reflect on how plant species adjust their gas exchange metabolism, interplay of CO_2_ and H_2_O acquisition and use, and adaptation patterns to different environments (Dawson et al. 2002). McKown et al. (2014a) reported a 6.6‰ range in Δ^13^C values among 461 natural accessions of *P. trichocarpa.* Often the variations in *WUE*_i_ and Δ^13^C are associated with the variations in photosynthetic capacity of the populations (Pointeau and Guy 2014). In this study, even though no linkages were observed between Δ^13^C and geo-climatic variables, we found trait associations between Δ^13^C, PNUE and leaf N. Such variations in Δ^13^C and their relative role in photosynthetic capacity and adaptation have been studied in many trees (Anderson et al. 1996; Monclus et al. 2005).

Intraspecific variation in nitrogen uptake and assimilation may differ among plant populations adapted to temperate (NO_3_ dominant soils) and boreal (NH_4_ dominant soils) climates. Soil derived NO_3_ nitrogen is assimilated by the nitrate reductase (NR) and nitrite reductase (NiR) pathway, producing NH_4_. Subsequently, soil derived NH_4_ along with NO_3_ derived NH_4_ is assimilated via the glutamine synthetase (GS) and ferredoxin glutamate synthase (fd-GOGAT) pathway resulting in δ^15^N variations of plant tissues (Lopes and Araus 2006). Hence, natural abundance of δ^15^N in a plant provides an insight into the causal relationships between uptake, assimilation and allocation of nitrogen (Kalcsits and Guy, 2013). If NO_3_ is partially assimilated in roots than shoots are enriched in δ^15^N or if wholly assimilated in roots or shoots than shoots are not enriched in δ^15^N. We observed between 2 and 4‰ within plant variation (leaf *vs*. wood), and this could be due to partial assimilation of source nitrogen (particularly, NO_3_) in the roots, resulting in isotopic differences between tissue types (Evans et al. 1996). The observed latitudinal clines in δ^15^N imply that there is an adaptive genetic variation in assimilation of NO_3_ nitrogen between roots and shoots in *S.eriocephala*.

### Geographic variation in *Salix* seasonality

Functional traits that explain ecophysiological capacities are constantly modified during the growing season as a result of growth cessation and bud set (McKown et al. 2013). For instance, bud set has high heritability across multiple years (*H*^2^=0.739), whereas vegetative traits such as, leaf mass per unit area (*H*^2^=0.810 spring; *H*^2^=0.150 post bud set) is more plastic within a given season (McKown et al. 2014a). A number of common garden studies under single photoperiodic regime suggest daylength sensitivity in bud phenology to have a genetic basis (Ingvarsson et al. 2006, McKown et al. 2014a). For most deciduous trees, having met the chilling needs (endo-dormancy), the onset of spring bud flush marks the shift from a dormant,restive stage (eco-dormancy) to an active growth stage upon accumulation of necessary heat sums under favourable environmental conditions (Worrall 1993).

Spring leaf emergence has been shown to have advanced over the past century with a steady lengthening of growing seasons (McMahon et al. 2010) leading to increased carbon fixation by terrestrial plants (Peñuelas et al. 2009). On the other hand, this increase in carbon sequestration is partially offset by enhanced rates of respiration (Piao et al. 2008). Temperature driven spring green-up often displays lower broad-sense heritability (*H*^2^=0.43 to 0.68, Tsarouhas et al. 2003) as temperatures fluctuate a lot from year-to-year. Previous common garden studies have reported a narrow range for bud flush (∼1-3 weeks) among intraspecific populations that display a latitudinal cline (*Acer saccharum* Marsh., Kriebel 1957; *Betula alleghaniensis*, Clausen and Garrett 1969; *P.balsamifera* and *P. tremula* L., Soolanayakanahally et al. 2013, 2015; *P. trichocarpa*, McKown et al. 2014a), but higher spring temperature can shorten the duration for bud flush. So, under common garden environments, trees from low latitudes often display later bud flush due to higher chilling and heat unit needs than the trees from high latitudes (Hannerz et al. 2003). Weih (2009) study emphasis spring leaf emergence and leaf abscission event’s being critical for biomass accumulation by *Salix* species and it is important to determine the impacts of future spring temperature change on the timing of bud flush at a given latitude.

While *Populus* has been a focus of extensive works in understanding the molecular mechanisms of autumn phenology primarily cued by photoperiod (Ingvarsson et al. 2006, Keller et al. 2011) such an understanding is lacking for *Salix* (Hallingbäck et al. 2015). The seasonal variation in photoperiod is consistent from year-to-year and is a reliable cue for onset of bud set, leaf senescence and induction of dormancy than temperature which is far less predictable and shows seasonal fluctuations (Barr et al. 2004). Our observed latitudinal cline in the onset of senescence and leaf drop is consistent with our a *priori* expectations. As willow plants attain competency to respond to photoperiod by mid-summer, they would have to wait for the critical daylength to induce autumn phenological events (Soolanayakanahally et al. 2013), with high latitude genotypes ceasing growth under longer critical daylength than the trees from low latitude (Pauley and Perry 1954, Howe et al. 1995).

At high latitudes, greater susceptibility to insect and disease is largely explained by evolution of plant defenses which display latitudinal clines (Anstett et al. 2015). As observed in this study, geographic regions that experience low *Melampsora* rust occurrence, natural resistance could be negatively selected in the absence of biotic stressors. In addition, larger stomatal pore length among high-latitude *S.eriocephala* genotypes might provide greater surface area for *Melampsora* rust spores to penetrate and colonise a given leaf surface area compared to low-latitudegenotypes. Our results find support for the hypothesis“*carbon gain and disease resistance trade-offs*” by McKown et al. (2014b), the notion that fast growth might have negative fitness with disease resistance. In addition, an inherent resistance to *Melampsora* rust is metabolically costly with substantial increase in certain classes of metabolites, particularly, tannins.

### 4.4 Adaptive phenotypic trait convergence within *Salicaceae*

In the Northern Hemisphere the members of Salicaceae (*P.balsamifera* and *S. eriocephala*) are sympatric species with overlapping natural ranges across north temperate-boreal climates (Hosie 1979, Dorn 1970). Both are restricted to moist and nutrient rich sites, exhibit indeterminate growth, and have a much wider north-south range. In general, parallel evolutionary selection pressures produce functionally convergent phenotypic traits in related taxa. Broadly, we hypothesised that the patterns in form and function convergence on a similar adaptive phenotype among *Populus* and *Salix* species in spite their long divergence (∼65 million years).

In a greenhouse study, without any resource limitations *P.balsamifera* populations displayed latitudinal gradients in photosynthesis (*A*), whereby higher *A* was ably supported by enhanced *g*_m_ and leaf N at high latitudes (Soolanayakanahally et al. 2009). Similar mechanism in *S.eriocephala* was observed in an outdoor common garden, whereby higher *A* in genotypes from high latitudes is partly mediated by higher *g*_m_ and *g*_s_ (mediated by larger stomatal pore length rather than higher density). These two studies collectively demonstrate a very strong correlation between *A* and latitude of origins and thus suggest that the possibility of a convergent adaptive phenotypic trait selection to compensate for short growing seasons. In addition, both species were hypostomatous with stomatal density displaying a strong negative association with latitude of origin. Unlike greenhouse grown *P. balsamifera*, *g*_s_ was positivelycorrelated with latitude of origin in *S.eriocephala*. In addition, Leaf N, LMA and CCI were positively associated with each other contributing to greater *A* at high latitudes in both species. Four independent common garden studies along a latitudinal gradient by the same group (Soolanayakanahally et al. 2009, 2015, McKown et al. 2014a, present study) and an additional study by Kaluthota et al. (2015) showed that genetic divergence in *Salicaceae* members largely explains the variation observed in functionally important leaf traits-*A*, LMA, Leaf N.

Further, both species display a strong latitudinal cline in autumn phenology (leaf senescence, green cover period, leaf drop) resulting from photoperiodic adaptation (Soolanayakanahally et al. 2013). Overall, our findings in *S.eriocephala* and *P.balsamifera* lend support to the hypothesis that natural selective pressures enacted along similar environmental gradients led to phenotypic trait convergence in sympatric *Salicaceae* members.

## 5. CONCLUSION

Our common garden results speak to the paramount role of adaptive trade-offs along latitudinal gradients, suggesting that certain trait combinations have been favoured by natural selection, leading to a locally adapted phenotype. First, we found multiple evidences for an enhanced photosynthetic assimilation rate (*A*) at high latitudes ably supported by stomatal traits (increased *g*_s_, and larger stomatal pore length) and a greater *g*_m_ which all coevolved along geo-climatic gradients. In addition, higher *A* at high latitudes results from greater LMA with higher leaf N concentrations. Taken together, our results highlight latitudinal clines in *A* as an adaptation to growing season length. Second, we observed least variations in water use-efficiency as determined by Δ^13^C values among genotypes from different latitudes with varying precipitation patterns. Observed latitudinal cline in δ^15^N values suggests that NO_3_ nitrogen is partially assimilated inthe roots leading to enrichment of stem wood tissue. Last, a strong photoperiodic adaption was observed in autumn phenology traits accounting for high heritability that could be exploited in willow improvement program for biomass and environmental applications. Overall, an adaptive negative relationship between stomatal density and pore length is optimised to achieve maximum leaf diffusive conductance to CO_2_ within the physiological framework of carbon/water balance across a range of a latitudes and climates.

## AUTHOR CONTRIBUTION

A.S.K.S. participated in data analysis and interpretation and drafted the manuscript. R.Y.S. conceived the study, performed gas exchange measurements, participated in analysis and interpretation, and edited the manuscript. R.D.G. interpreted the stable isotope results and complemented the writing. The authors declare that the research was conducted in the absence of any commercial or financial relationships that could be construed as a potential conflict of interest.

## ACKNOWLEDGMENTS

The efforts of Don Reynard and Chris Stefner in establishing the willow common garden is wholly appreciated. We thank Hamid Naeem for monitoring phenology over two seasons. This work was funded to R.Y.S. by the Agriculture and Agri-Food Canada (LOI 1268) and by the Natural Sciences and Engineering Research Council of Canada (NSERC) Discovery Grants Program to R.D.G.A.S.K.S. is a NSERC Visiting Fellow at AAFC. The authors express thanks to George Argus for assistance with willow specimen identification.

## Supplementary Table

**Table ST1.**
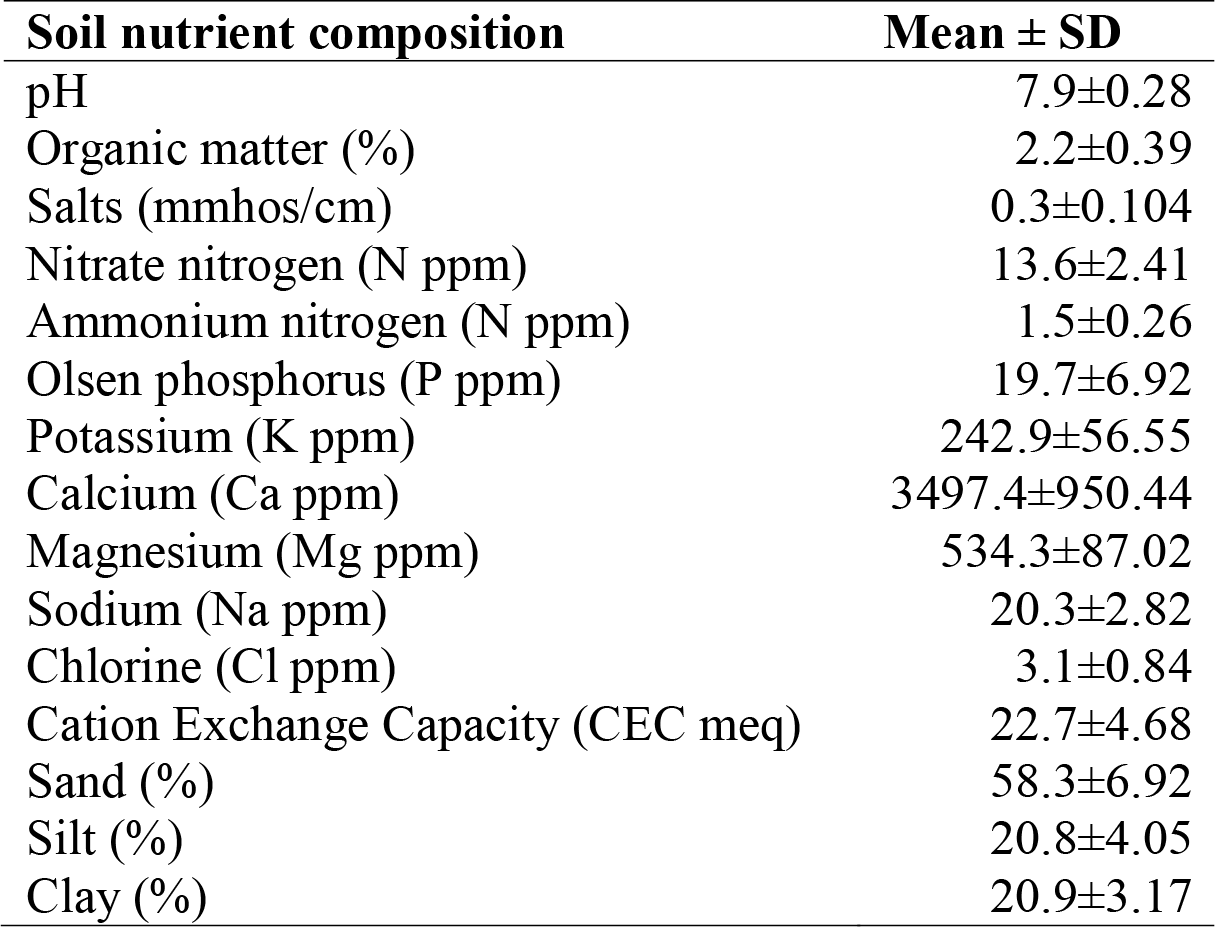
Soil analysis results from 15cm depth at Indian Head common garden, Saskatchewan in 2012. Values shown are means ± SD of nine individual samples analyzed from sandy clay loam texture soil.

## Supplementary Figure

**Figure SF1A.**
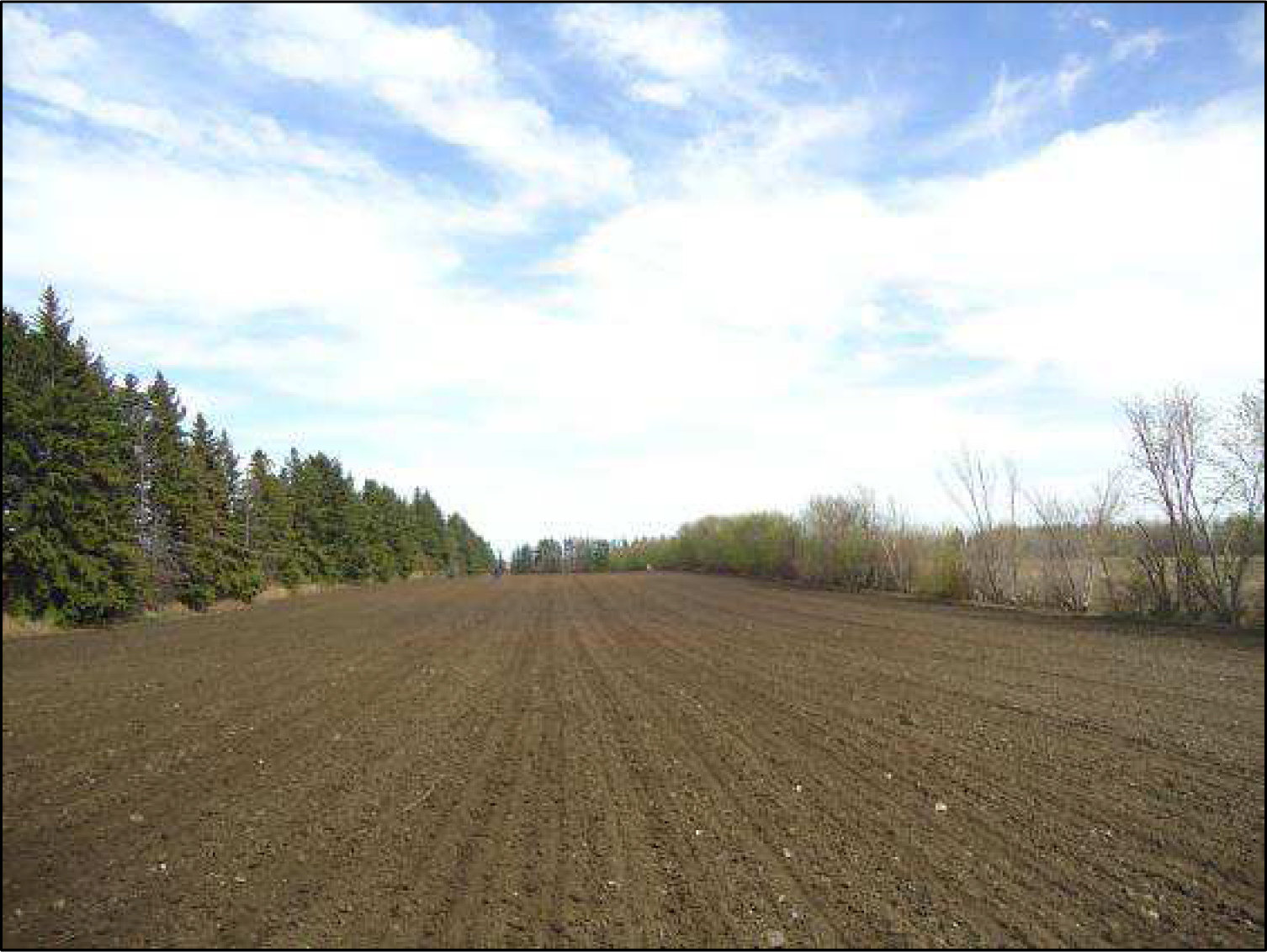
Pictorial description of site preparation, planting, management and trait measurements in the common garden at Indian Head research station, Canada Common garden site preparation prior to willow planting

**Figure SF1B.**
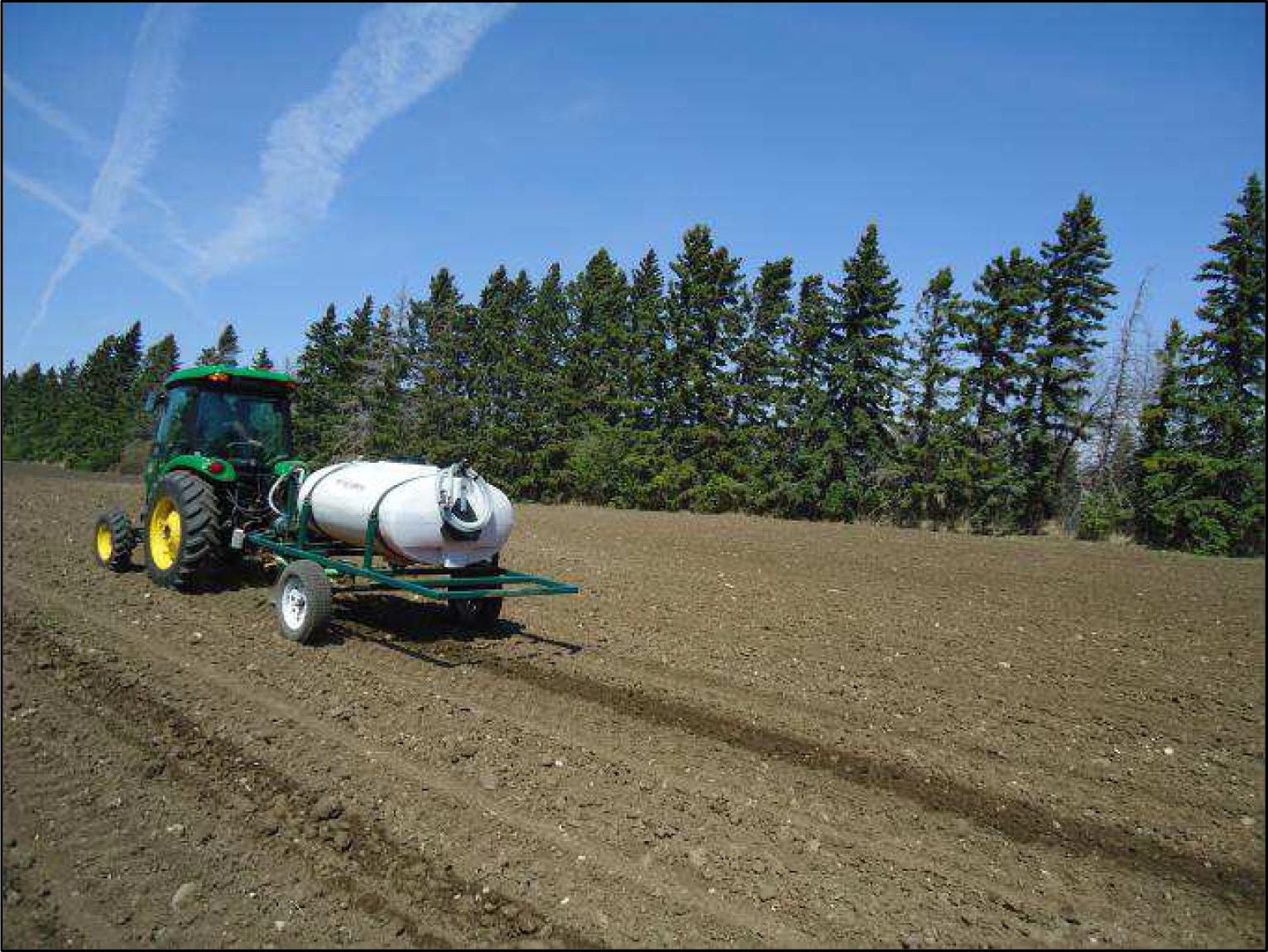
Nutrient amendments to sandy clay loam soils along planting strips.

**Figure SF1C.**
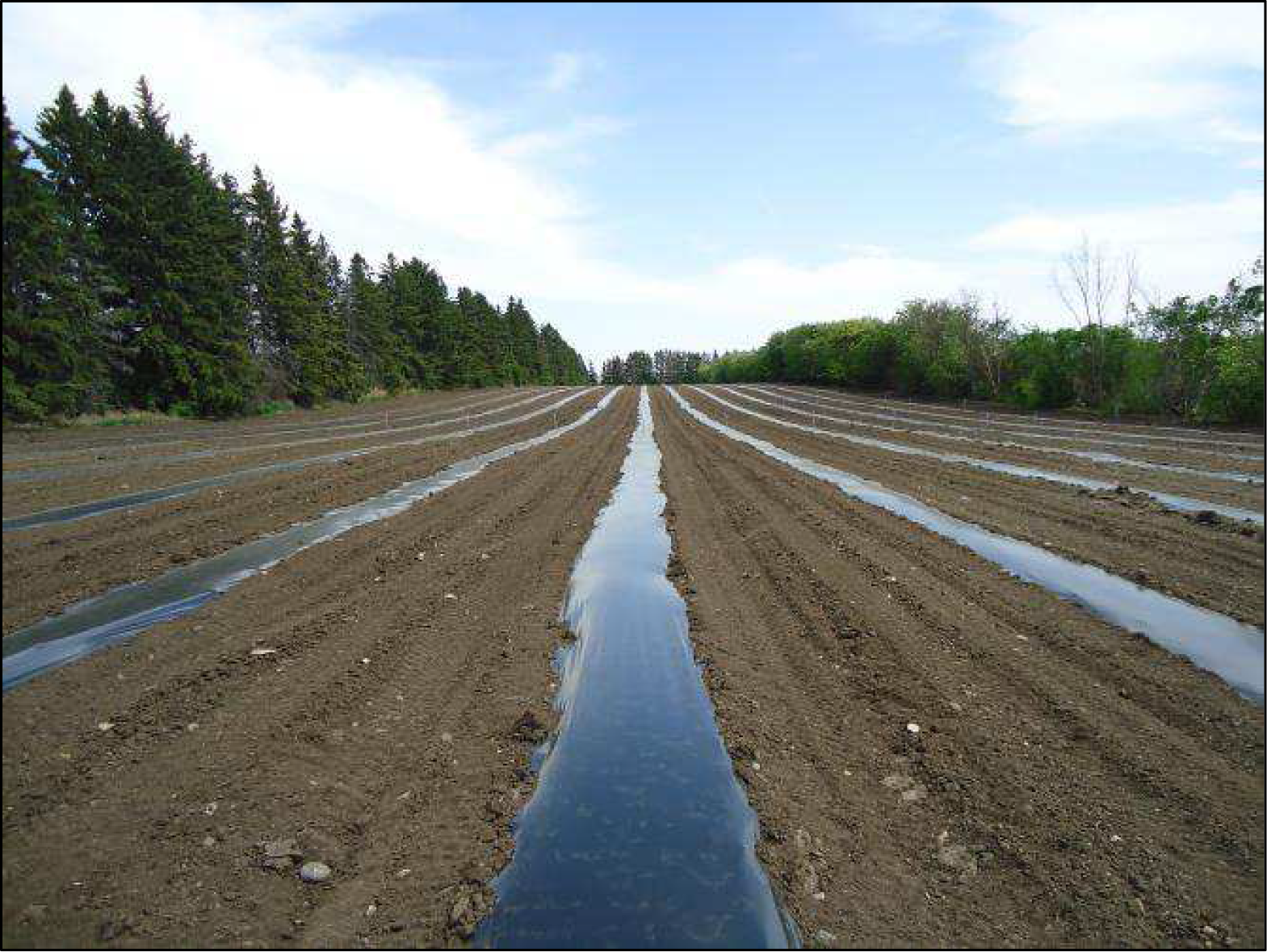
Laying of plastic mulch to avoid intra row weed competition.

**Figure SF1D.**
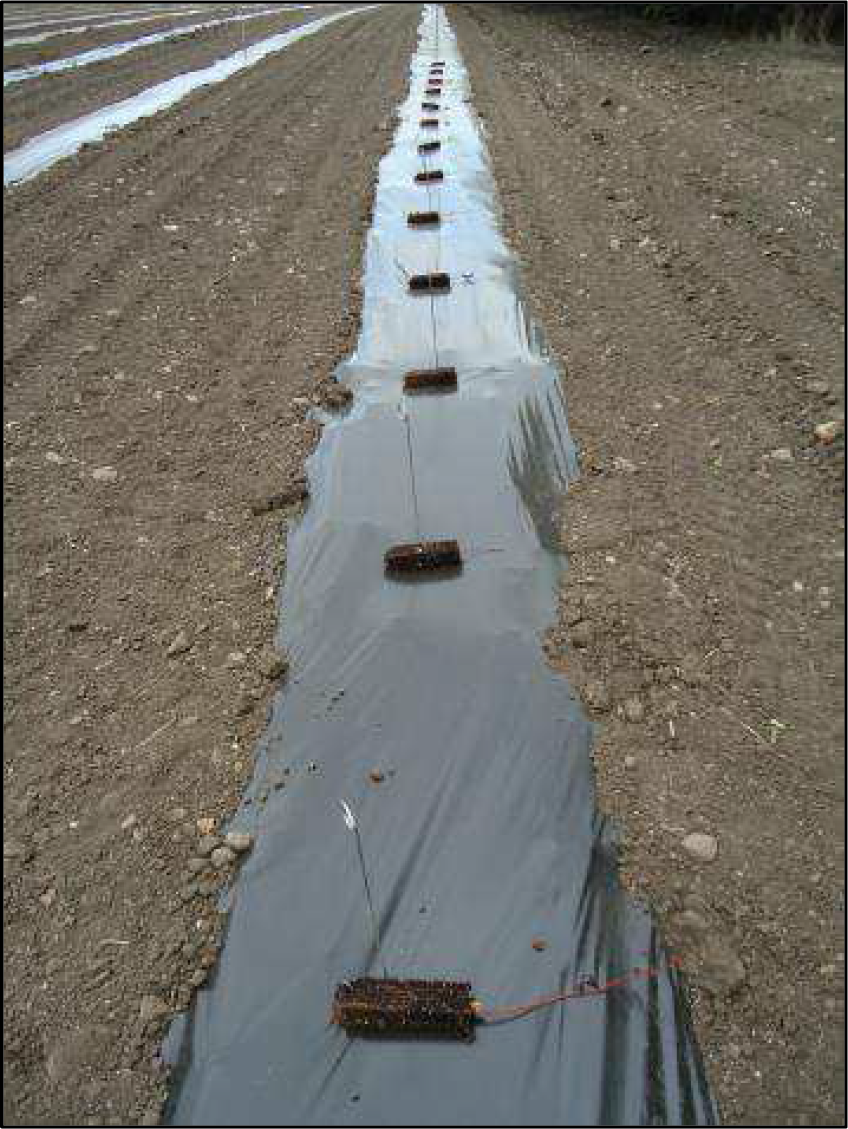
Dormant root plugs ready for planting at one metre intervals during the spring of 2012

**Figure SF1E.**
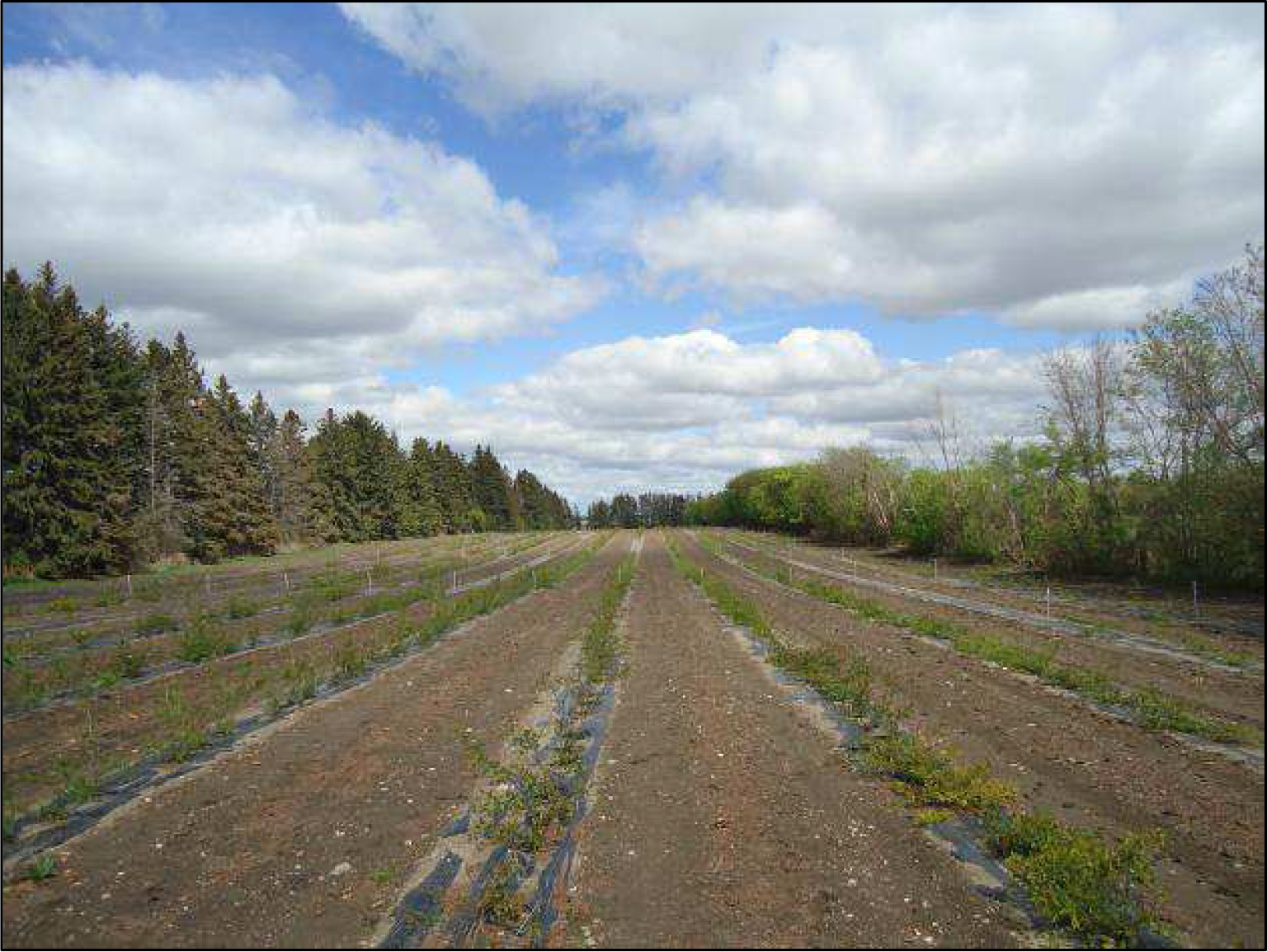
Uniform establishment of willow common garden during the summer of 2012

**Figure SF1F.**
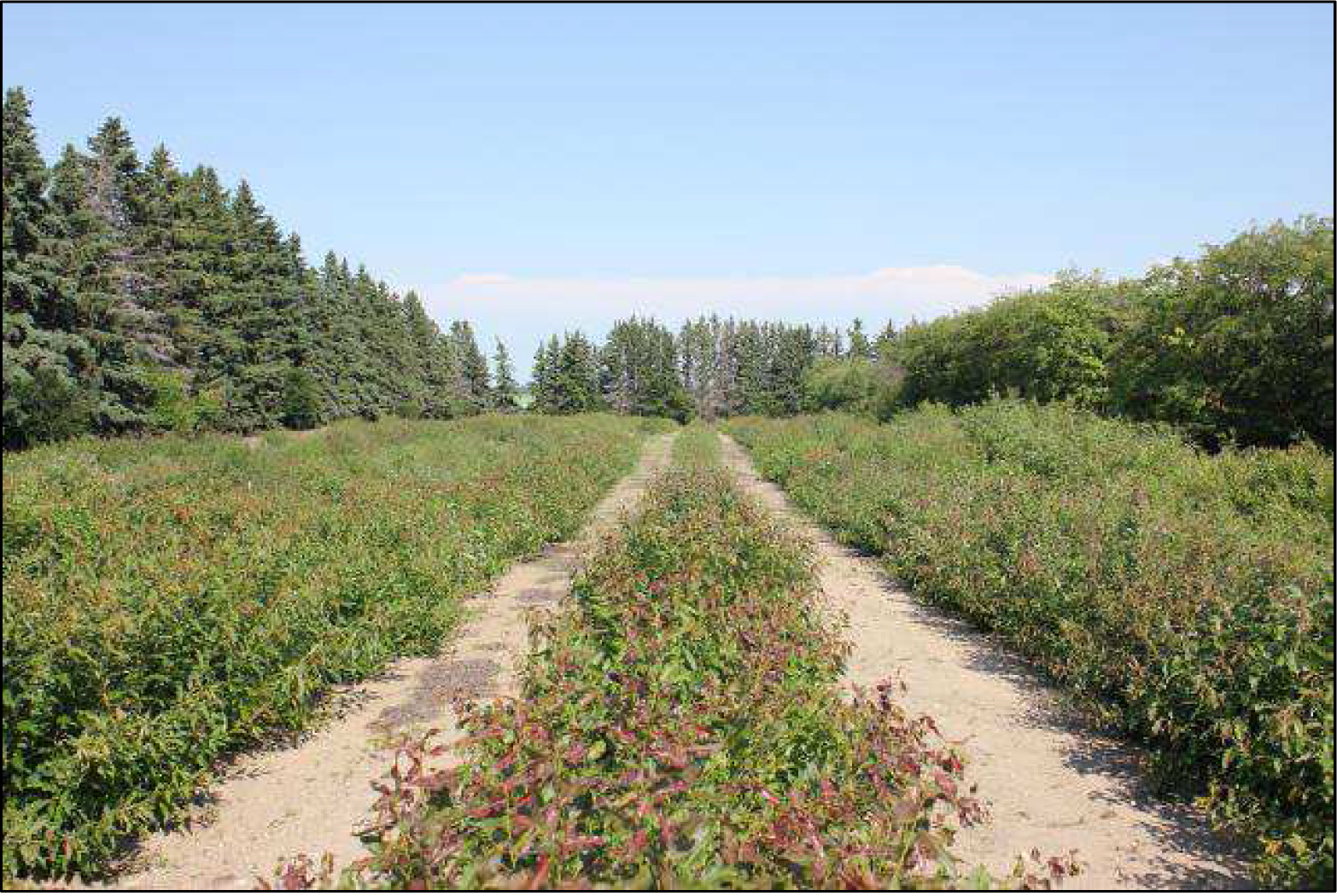
Willow growth during the summer of 2014.

**Figure SF1G.**
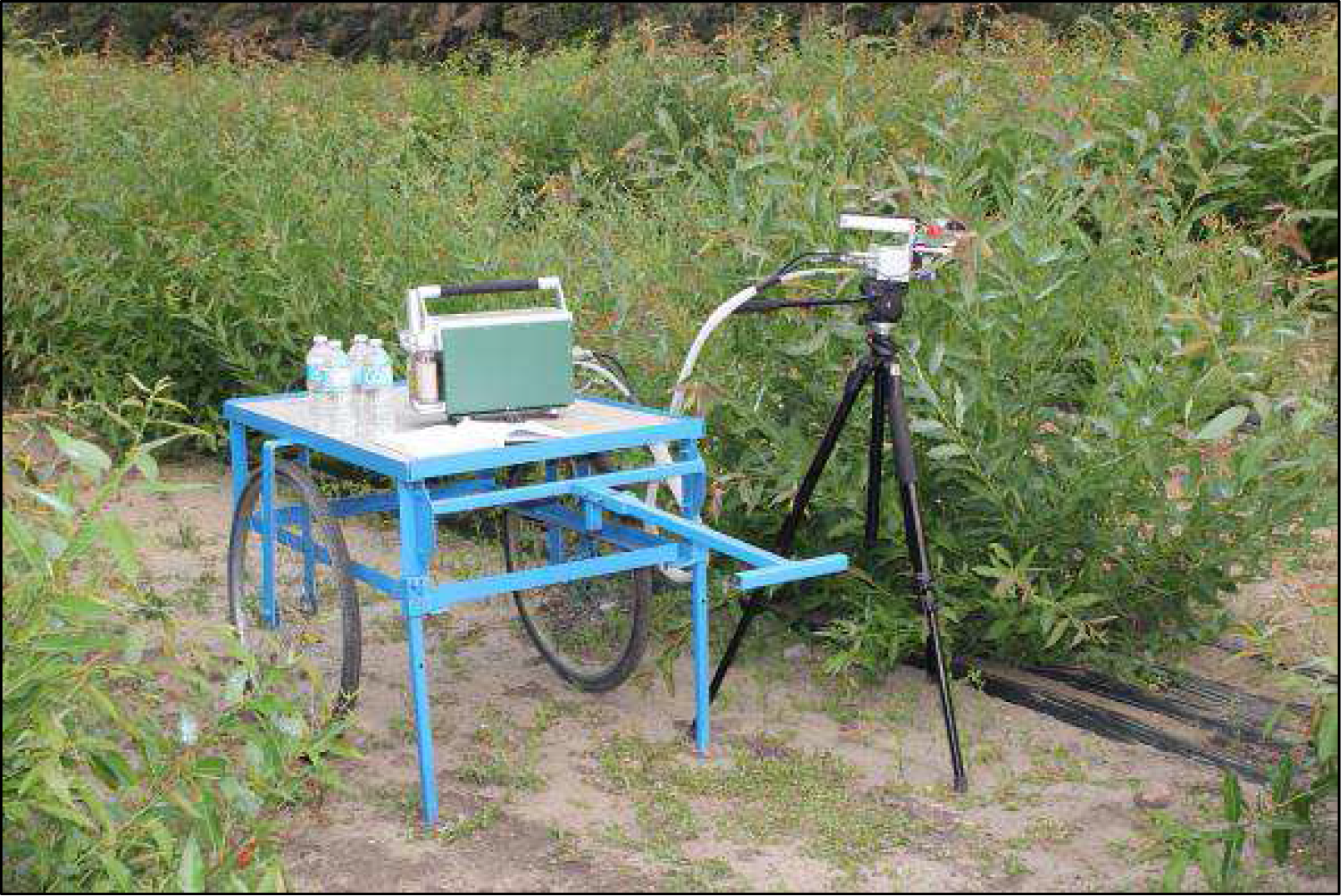
Gas exchange measurement during active growth in the summer of 2014.

